# Cognitive and autonomic physiological responses to daily transcutaneous auricular vagus nerve stimulation in healthy adults

**DOI:** 10.64898/2025.12.12.693915

**Authors:** Shawn D. Flanagan, Rodman D. Tompkins, Juliana R. Hougland, Meaghan E. Hazelet, Danielle Moss, Chris Connaboy

## Abstract

**Introduction:** The vagus nerve regulates autonomic processes central to cognition. Transcutaneous auricular vagus nerve stimulation (taVNS) can acutely modulate executive function in healthy adults, but the effects of repeated dosing, roles of executive subdomains, and importance of sex as a biological variable require clarification.

**Method:** In a randomized, single-blind, sham-controlled, parallel group study, 12 healthy adults (six women) received 10 consecutive days of left monoaural taVNS (VNS) or sham (CON), with a two-week follow-up rechallenge. Stimulation (100Hz; 200µs pulse width) was delivered continuously (online) at rest and during the Eriksen Flanker (inhibition), Wisconsin Card Sorting (shifting), and Sternberg (working memory) tasks. Resting autonomic measures included pupil diameter, skin conductance (SCL), mean R-R interval (mRR), respiration, and blood pressure. Primary outcomes were analyzed with linear mixed models and Šidák-corrected contrasts.

**Results:** Ten days of taVNS produced immediate and cumulative changes in multiple executive function subdomains that remained evident at rechallenge, with the strongest effects on accuracy. Across tasks, benefits were most pronounced in women; men displayed initial performance costs, but with greater longitudinal improvements. Cognitive effects were most evident on higher-difficulty trials. Autonomically, taVNS increased SCL during and after stimulation, amplified stimulus- and task-evoked pupil responses, and transiently lengthened mRR. Stimulation was well tolerated and blinding was acceptable.

**Conclusions:** Resolution of taVNS effects to higher-difficulty trials supports adaptive-gain modulation within executive function networks. The combination of larger evoked pupil/SCR, transient mRR lengthening, and increased SCL indicates that taVNS increased autonomic flexibility and arousal. Our findings identify biological sex and stimulation intensity as potentially important considerations for taVNS efficacy and provide longitudinal estimates to guide dosing, biomarker selection, and trial design. Collectively, the results support continued development of taVNS as a safe tool for the modulation of executive function and autonomic activity in healthy adults.

## INTRODUCTION

Transcutaneous auricular vagus nerve stimulation (taVNS) is a noninvasive brain stimulation technique that has garnered significant interest for its potential to improve health and performance. By delivering weak electrical current to Aβ afferents of the auricular branch of the vagus nerve (ABVN) in the outer ear, taVNS seeks to reproduce the central effects of invasive VNS without the risks, cost, or burdens of surgical implantation [1-5]. The bottom-up effects of taVNS are thought to arise primarily from the activation of *nucleus tractus solitarius* (NTS) in the dorsal medulla [6-8]. From NTS, second- and third-order projections engage nuclei within the central autonomic network (CAN) and reticular activating system (RAS), influencing monoaminergic, GABAergic, cholinergic, and glutamatergic signaling across subcortical and cortical nodes of the frontoparietal (FPN), salience (SN), limbic (LN), and default mode (DMN) networks [9-13]. Through these circuits, taVNS can modulate attention, learning, stress reactivity, motor behavior, immune function, and core homeostatic processes that contribute to neuropsychiatric and autonomic disorders when dysregulated [14-18]. Beyond therapeutic contexts, taVNS is increasingly used to augment cognitive and autonomic functions in healthy adults [19-22].

A central premise of neuromodulation is that durable benefits arise from the consolidation of adaptive responses to *repeated dosing* [11, 23, 24]. In healthy adults, acute taVNS yields small-to-moderate improvements in cognition overall (meta-analytic *g* = 0.21), with the most consistent gains in executive function (meta-analytic *g* = 0.66) despite heterogenous effects on core subdomains such as inhibition, shifting, and working memory (WM) [25-29]. These effects are commonly attributed to the modulation of locus coeruleus-norepinephrine (LC-NE), basal forebrain-cholinergic (BFCN), and GABAergic signaling within FPN and SN [11, 30, 31]. As indirect indices of central arousal and autonomic control, pupil diameter and cardiorespiratory activity are frequently used as mechanistic proxies of acute taVNS neuromodulatory effects on LC-NE, BFCN, and GABAergic signaling, although their specificity to any single neuromodulatory system is limited and protocol-dependent [32-34].

In contrast to single-session studies, the consequences of repeated (i.e., daily) taVNS remain unclear, including whether effects accumulate and persist at follow-up rechallenge. In addition, while sex differences in executive function are debated and generally small compared with other factors [35], women appear more responsive to certain therapeutic neuromodulation interventions such as repetitive transcranial magnetic stimulation [36]. Potential contributors include variation in endogenous or contraceptive-related neurosteroid concentrations [37-39] and structural differences within FPN/SN regions implicated in executive function and taVNS cognitive effects [40, 41]. Sex-specific responses to acute taVNS have been reported [42-46], but whether daily taVNS elicits sexually dimorphic changes in executive function or autonomic physiology in healthy adults is largely untested.

In this randomized, single-blind, sham-controlled, parallel group study, we tested whether daily monoaural taVNS modulates executive function and autonomic physiology across 10 consecutive laboratory sessions and at a two-week follow-up rechallenge. To dissociate effects across core executive function subdomains, we used three well-characterized tasks: Eriksen Flanker (inhibition), Wisconsin Card Sorting (shifting), and Sternberg Working Memory (WM). To probe potential mechanisms, we quantified resting and evoked autonomic biomarkers (pupil diameter, skin conductance, cardiorespiratory activity) with demonstrated, though inconsistent, sensitivity to acute taVNS. Our aims were to: 1) quantify change with repeated dosing and its persistence; 2) resolve effects within executive function subdomains; and 3) explore sex as a biological variable. Based on single-session reports, we hypothesized that repeated taVNS would enhance shifting and WM accuracy and modulate physiological indices consistent with LC-NE engagement, with rechallenge effects detectable at follow-up. Using dense repeated sampling under tightly controlled conditions, this exploratory study was designed to address key evidence gaps and to provide estimates for parameter selection and hypothesis specification in confirmatory trials.

## MATERIALS AND METHODS

### Participants

Twelve healthy young adults (*n* = 6 women; age, 25.6 ± 4.4 yr; height, 172.3 ± 7.7 cm; weight, 71.3 ± 9.3 kg; handedness, 93.2 ± 12.3 LQ [47]; physical activity, 90.4 ± 59.6 MET hr . wk^-1^ [48]) were enrolled as a university convenience sample (**Table 1**). All participants were right-handed and reported normal sensory and cognitive status and normal or corrected-to-normal vision. Exclusion criteria included: history of adverse reactions to taVNS; pregnancy; immunocompromised status; hemophilia; psoriasis vulgaris at electrode sites; vagal hypersensitivity; current neurologic, psychiatric, or other mental-health disorder; psychoactive medication use; and history of alcohol or substance abuse. All provided written informed consent. The study was approved by the Rosalind Franklin University Institutional Review Board (SCPM24-429) and Air Force Human Research Protection Office (FWR20240060X).

**Table 1.** Participant Characteristics.

| Group | VNS |  |  | CON |  |  |
| --- | --- | --- | --- | --- | --- | --- |
| Age (yr) | 26.8 | ± | 5.9 | 24.3 | ± | 1.8 |
| Height (cm) | 172.5 | ± | 7.5 | 172.2 | ± | 8.7 |
| Body mass (kg) | 68.7 | ± | 11.1 | 73.9 | ± | 7.2 |
| Handedness (LQ) | 94.8 | ± | 10.0 | 91.7 | ± | 15.1 |
| IPAQ (MET hr · wk<sup>-1</sup>) | 56.9 | ± | 18.6 | 123.9 | ± | 69.1 |

Values are means ± SD. Independent unequal variance t-tests confirmed similar age, height, body mass, handedness, and physical activity between groups ( $p > 0.05$ ). IPAQ, International Physical Activity Questionnaire.

### Experimental Approach

A randomized, sham-controlled, single-blind (participants), parallel group design was used to assess immediate and cumulative cognitive and autonomic physiological responses to daily taVNS along with rechallenge effects after a two-week intersession interval. Following consent and familiarization (including task practice), participants were randomly assigned (computer-generated; sex-balanced) to active taVNS (VNS; *n* = 6, 3 women) or sham (CON; *n* = 6, 3 women) and received stimulation once daily for 10 consecutive days (**Fig 1A/B**). At the two-week follow-up (F), participants underwent an on-stimulation rechallenge with identical parameters to assess long-interval responsiveness. These healthy adults were asked to maintain stable diet, exercise, and medication routines during the study. Questionnaires on sleep quantity/quality [49], muscle soreness [50], and life stress [51] were completed at the beginning of each visit. Given that hydration can influence cognition [52, 53], urine specific gravity (USG; Master URC, Atago) was measured at the start of each visit. In the event of dehydration (USG ≥ 1.025), water was provided before procedures and *ad libitum* thereafter. Hydration, sleep, soreness, and stress were comparable between groups and visits (**Table S1**). To reduce diurnal variability, visits were completed at similar times for each subject (± 2 hr).

**Figure 1.**
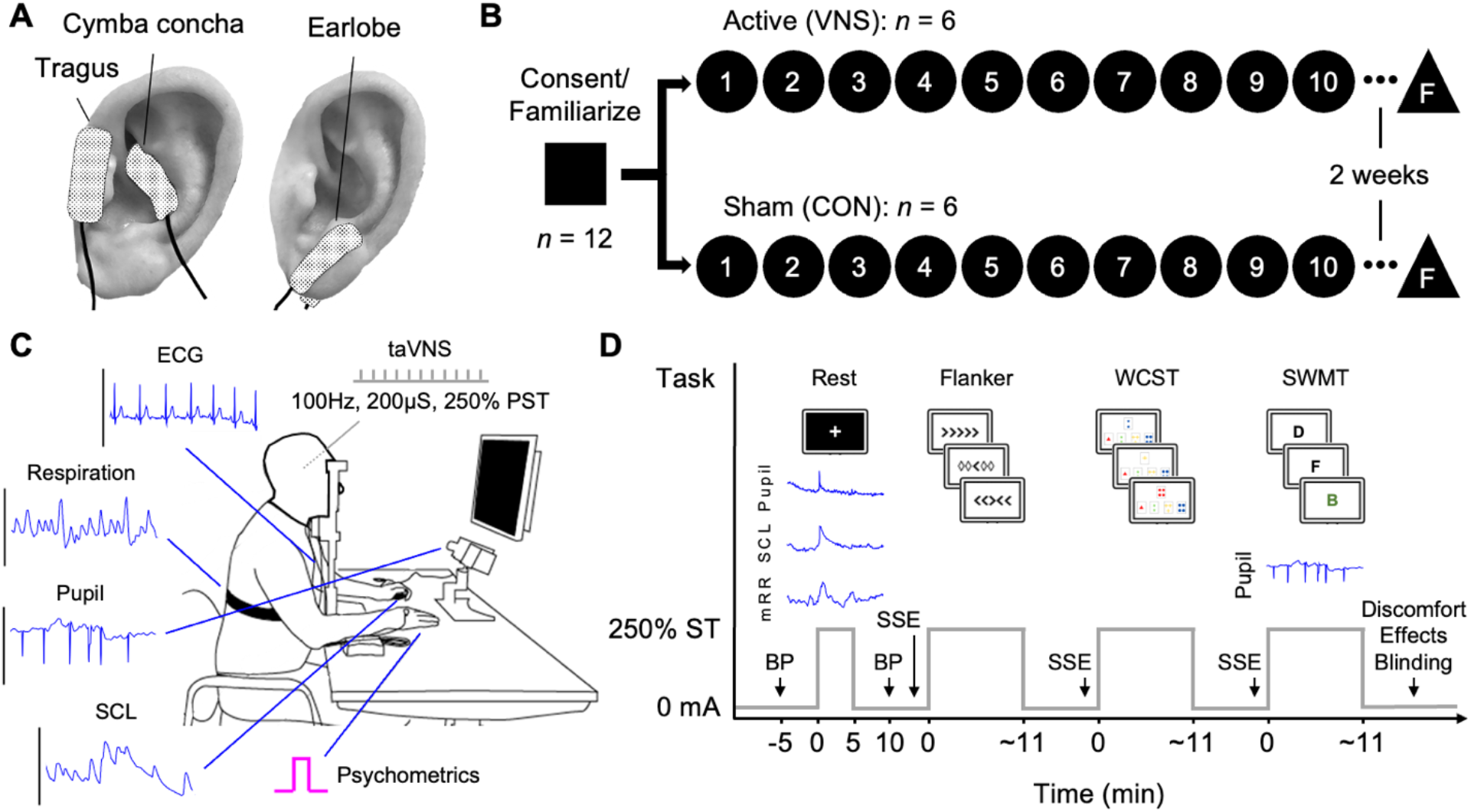
Experimental Approach. **A)** Electrode placement for active (VNS) and sham (CON) transcutaneous auricular vagus nerve stimulation (taVNS). **B)** The study used a randomized, single-blind, sham-controlled, parallel group design to assess whether daily monoaural taVNS modulates cognitive performance and autonomic physiological activity in healthy young adults across 10 consecutive laboratory sessions and at a two-week rechallenge follow-up (F) visit (*n* = 12, sex-balanced). **C)** Study procedures were performed in a quiet room with consistent ambient lighting and the head stabilized in a desk-mounted chin/forehead rest. Continuous stimulation was applied at 250% perceptual sensory threshold (PST) and 100 Hz with a 200 µs pulse width. Measurements included pupillometry, skin conductance level (SCL), ECG, respiration, blood pressure (BP), and psychometrics (response times, errors, lapses). **D)** Each visit began with assessment of resting autonomic physiological responses to taVNS based on comparisons among contiguous 5-min baseline (Pre), stimulation (Stim), and post-stimulation (Post) segments. Next, the Eriksen Flanker Task (Flanker; N = 120 trials), Wisconsin Card Sorting Test (WCST; N = 120 trials), and Sternberg Working Memory Task (SWMT, N = 60 trials) were performed with stimulation delivered from the beginning to end of each task (∼11min each). Pupil diameter was measured during each SWMT memorization and recall phase. The situational self-efficacy questionnaire (SSE) was completed before each cognitive task. At visit end, discomforts, perceived effects, and predicted treatment condition were assessed.

Each experimental visit began with confirmation of study controls, sensor placement, device calibrations, and perceptual sensory threshold measurements. Pupillometry, skin conductance level (SCL), ECG, and respiration were recorded during a 5-min baseline, 5-min of continuous stimulation (Stim), and a 5-min recovery period (Post). During this resting assessment, participants were seated and fixated a central cross at eye level (**Fig 1C/D**). Blood pressure (BP) was measured immediately before and after the assessment. Continuous stimulation was then applied during three tests of executive function (Eriksen Flanker Task [Flanker], Wisconsin Card Sorting Test [WCST], and Sternberg Working Memory Task [SWMT]). At visit end, participants reported any discomforts or perceived effects and guessed their group allocation [54]; blinding success was summarized with Bang’s Blinding Index [55]. All measurements were obtained in a quiet room with consistent ambient lighting and with the head stabilized in a desk-mounted chin/forehead rest (SR Research).

### Transcutaneous Auricular Vagus Nerve Stimulation

Active and sham taVNS were delivered to the left ear with a constant-current stimulator (DS7R, Digitimer). In accordance with parametric optimization studies for taVNS-induced NTS/LC fMRI and autonomic physiological activity [44, 56], stimulation was delivered at 100 Hz with 200 µS monophasic rectangular pulses at 250% of each participant’s daily perceptual sensory threshold (0.1 mA staircase method with modified starting phase; thresholds: VNS = 1.5 ± 0.6 mA, CON = 1.9 ± 0.6 mA; intensities: VNS = 3.7 ± 1.5 mA, CON = 4.9 ± 1.6 mA). The parameters chosen for stimulation site, amplitude, duration, and timing were similar to those commonly used in studies on single session taVNS effects [25, 57, 58]. For VNS, the cathode was centered on the cymba concha (depolarization site) and the anode on the tragus (**Fig 1A**). For CON, the cathode (anode) was centered on the anterior (posterior) earlobe. Disposable 25 × 10 mm Ag/AgCl hydrogel electrodes (N306, Neotech) with hydrocolloid ends and PVC-jacketed lead wire (DIN 42802, 1.5 mm touch-proof) were used. Skin was cleaned with alcohol swabs before electrode placement, and the electrodes were reinforced with medical tape (Transpore, 3M) as needed.

### Cognitive Function

The executive function subdomains targeted in this investigation were selected based on demonstrated behavioral sensitivity to taVNS and shared neuroanatomical correlates (e.g., FPN and SN engagement) [9, 11, 59-64]. The tasks (E-Prime 3.0, PST) were presented on a 17″ monitor (1280 × 1024 at 60 Hz) at eye level and a standardized viewing distance (76.2 cm). Participants were instructed to respond as quickly and accurately as possible. Stimulation was delivered continuously from the start to end of each task (i.e., online), with similar durations across groups, sexes, and visits (Flanker 666.1 ± 143.5 s; WCST 670.4 ± 143.4 s; SWMT 678.8 ± 143.2 s; **Table S2**). To minimize the potential influence of initial learning effects, each task was practiced to proficiency during the familiarization visit. In addition, to account for potential differences in self-efficacy, participants completed the situational self-efficacy (SSE) questionnaire before each task [65]. SSE scores did not differ between groups (Flanker, WCST, SWMT: *p* = 0.6936, 0.2236, 0.4692).

#### Eriksen Flanker Task

During the Flanker task, the goal is to respond to central (target) stimuli while ignoring adjacent (flanker) stimuli. The Flanker emphasizes response inhibition [66], an executive function subdomain behaviorally responsive to taVNS [27, 29, 67, 68]. Stimuli included five symbols (arrows or rhombuses), with difficulty progressively increasing from congruent (C) to neutral (N) to incongruent (I); participants indicated the direction of the central arrow with a left or right thumb response. Each visit included 120 trials (40 per type; total N = 15,840).

#### Wisconsin Card Sorting Test (WCST)

During the WCST, the goal is to correctly select cards in accordance with an unknown rule that changes without warning. The WCST is a complex task that emphasizes shifting [69], a subdomain with sensitivity to taVNS [25, 28]. Using a handheld mouse-controlled screen cursor, a response card was matched to one of four multidimensional stimulus cards according to a hidden sorting rule (symbol color, number, shape) inferred via trial-by-trial feedback. The sorting rule changed every 10 cards (one category). Each visit included 120 trials (3 categories × 10 trials × 4 replications; randomized; total N = 15,840).

#### Sternberg Working Memory Task (SWMT)

The goal of the SWMT is to correctly memorize and recall a specific list of letters. The SWMT principally evaluates short-term WM [70], a subdomain that is behaviorally responsive to taVNS [19, 21, 30]. An 8000 ms memorization (M) phase presented eight letters (1000 ms each), some black (memorize) and some green (ignore), followed by a recall (R) phase requiring yes/no responses to probes. Participants completed two sets of three blocks that varied by difficulty based on the number of letters to ignore: low (Lo) = 5/8, moderate (Mo) = 3/8, high (Hi) = 1/8.

Block order was randomized, with 60 trials per visit (N = 7,920). As an indirect physiological index of LC-NE/BFCN activity and cognitive load, pupil size was recorded during each M and R phase [71-75]. Stimuli were presented on a constant-luminance background, and letter colors were isoluminant.

### Autonomic Physiological Activity

#### Pupillometry

Binocular pupil size was recorded at 1000 Hz (Eyelink 1000 Plus, SR Research). Eye tracking performance was calibrated (9-point) and validated (10-point) in accordance with manufacturer guidelines. Pupil data were converted from arbitrary units to mm via an artificial pupil standard (76.2 cm distance) and multipoint calibration (Labchart v8, AD Instruments).

#### Skin conductance levels

Bipolar 8 mm Ag/AgCl finger electrodes measured SCL using constant-voltage exosomatic AC excitation (22 mV_rms_ at 75 Hz), 1 Hz second-order low-pass filtering, 12-bit DAC high-pass filtering, full isolation, 2000 Hz digitization, and 40 μS input range (FE116 GSR Amp; MLT117F Electrodes, AD Instruments) [76]. Electrodes were filled with conductive paste (Ten20, Weaver and Co), placed on the volar proximal phalanges of the left index and middle fingers, and secured with Velcro. After 5-10 min for skin hydration/equilibration, the system was zeroed (open-circuit, then subject baseline) and re-zeroed immediately before data collection. Recordings occurred in a climate- and humidity-controlled laboratory. Participants wore the sensors for a total of one hour.

#### Electrocardiography

ECG signals were measured using a bipolar three-lead setup (RA, LA, LL) with 16-bit resolution (±100 mV range), > 1 GΩ differential input impedance, > 100 db CMRR, < 0.5 µV_rms_ noise, and 4000 V_rms_ isolation (FE231 Bio Amp, AD Instruments; Kendall Medi-trace 530 Electrodes, Cardinal Health). Signals were digitized at 2000 Hz and band-pass filtered from 0.1-1000 Hz before analysis.

#### Respiration and blood pressure

Respiration was measured with a piezoelectric respiration belt (xiphoid level; 16/35 Powerlab; MLT1132 Respiration Belt; AD Instruments). Signals were digitized at 2000 Hz (±10 V input range, 16-bit; 1 MΩ input, > 100 db CMRR, 1.5 LSB noise). Blood pressure (BP) was measured manually (right arm; two assessors) before and after the resting assessment with participants seated, feet flat, and back supported (DuraShock Aneroid, Welch Allyn).

### Data Analysis

#### Cognitive function

Psychometric endpoints included median response times, error counts, and lapse counts. Response times were tabulated after the removal of false starts (< 200 ms), lapses (per-participant ROUT, Q = 0.1% [77]), and errors. Errors and lapses were assessed individually and combined (rejected trials). For the WCST, errors/lapses were categorized as perseverative (P; responses that occurred after the first trial in each category and consecutive errors after the first P error) or non-perseverative (NP; first category trials; first trials of categories 2-12; any trial after a correct response following one or more P errors).

#### Autonomic physiological activity

Pre-processed time series data for pupil, SCL, ECG, and respiration were segmented into three 5-min segments (Pre, Stim, Post); excess data at the beginning of Pre and end of Post (30 s each, plus 10-30 s jitter during Pre) were used for windowing. Pupillometry preprocessing followed current guidelines [71, 78, 79]: values < 1 mm (blinks) were padded by 100 samples and linearly interpolated; binocular traces were averaged, visually inspected, downsampled to 20 Hz, and averaged per segment. SCL was downsampled to 2 Hz and baseline-corrected (initial-value subtraction) before segment means were computed. ECG was detrended (smoothing priors; λ = 500; cutoff 0.035 Hz) and missed/misplaced/ectopic beats were corrected individually (Kubios Premium v3.5) [80-82]. Time-varying mean R-R interval (mRR) was computed using a 30 s window and 1 s step. Mean respiration frequency was computed per segment. To assess the possibility of transient (phasic) responses to stimulation onset, individual pupil, SCL, and mRR time series data were downsampled (pupil = 20Hz; SCL = 5Hz; mRR = 1Hz), windowed (pupil = -2 to +6 s; SCL = -20 to +60 s; mRR = -60 to +180s), and baseline-corrected (pre-stimulus mean subtraction). Response magnitudes (post-stimulus peak minus pre-stimulus mean) and processed time series data were retained for statistical analysis. No physiological data were lost due to poor quality.

### Statistical Analysis

Data are means ± SD unless otherwise noted. Analyses used Prism 10 (GraphPad), SPSS 30 (IBM), and JASP 0.95 (JASP Team). Participant, protocol, and control variables were analyzed with Welch’s *t*-tests and one-, two-, or three-way ANOVAs with Greenhouse-Geisser sphericity correction for models with repeated measures. Primary cognitive and physiological endpoints were analyzed with linear mixed models (LMMs). We used a simple to complex strategy: fixed effects were entered together; random intercepts and slopes were then added as supported; followed by residual covariance structures. Age was included as a covariate (mean). To account for the potential influence of variation in stimulation intensity by group and sex, individual values were decomposed into between-subject *z*-scores (participant mean vs group mean: I*z*_*between*_) and within-subject *z*-scores (session intensity vs participant mean: I*z*_*within*_) and added to each optimized LMM as covariates with interactions (group and sex). LMM selection was based on the lowest overall information criteria score (BIC, AIC, and -2LL) with definite positive Hessian matrix and successful convergence (**Table S3**). Random effect necessity was confirmed via Wald z statistics (*p* ≤ 0.10). The final models used REML with Satterthwaite *df* (≤ 100 iterations; max 10 step-halvings) [83]. Random effect covariance used variance components (VC); within-subject covariance structures were selected on theoretical grounds and fit indices subject to computational feasibility. Pairwise comparisons for LMM and ANOVA results were made with Šidák correction for multiple comparisons. Results are reported as EMM-based MD [95% CI], *t* statistic (Satterthwaite *df*), and 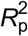 (LMM) or 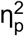 (ANOVA). Specific to stimulus-evoked physiological responses, in addition to LMMs (response magnitudes), time-series data were analyzed using factorial non-parametric permutation tests (Rde_kheradPajouh_renaud; 10,000 permutations) with cluster mass-based FWER correction [84, 85]. For this exploratory study, significance was set at *p* ≤ 0.10.

## RESULTS

Total daily stimulation time was 38.6 ± 7.2 min. Mean stimulation intensity was greater in CON than VNS (MD = 1.2 mA [0.0, 2.5], *t*(8) = 2.34, *p* = 0.0473,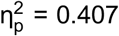) and in men than women (MD = 1.2 mA [-0.1, 2.4], *t*(8) = 2.19, *p* = 0.0598,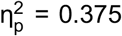), with no visit or interaction effects (**Table S4**). Associations and conditional contrasts for dose-response effects are provided in **S Results**. All LMM estimates below are evaluated at I*z*_*between*_ = 0 and I*z*_*within*_ = 0 (i.e., group/sex means and session means).

### Cognitive Function

#### Flanker

After removing 207 lapses (1.3% trials) and 362 errors (2.3%), 15,271 trials remained (**Fig 2A**). Median RTs differed by group across visits (group x visit: F_10, 77_ = 2.12, *p* = 0.0328), with an increase in RT at follow-up (F) in VNS compared with CON (MD = 31.2 ms [-0.9, 63.3], *t*(21.62) = 2.01, *p*_adj_ = 0.0569, 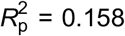 **Fig 2B**). Error and lapse rates differed by group and sex (group × sex: F_1, 22_ = 20.22, *p* = 0.0002; **Fig 2C**). Men in VNS had more rejected trials than men in CON (MD = 2.1 [0.9, 3.3], *t*(21.82) = 3.54, *p*_adj_ = 0.0018, 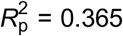 **Fig 2D**) and women in VNS (MD = 3.8 [1.9, 5.8], *t*(21.57) = 4.18, *p*_adj_ = 0.0004,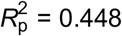). Conversely, women in VNS had fewer rejected trials than women in CON (MD = - 2.7 [-4.4, -1.1], *t*(21.62) = -3.40, *p*_ad_ = 0.0026, 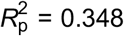). Given decreases in errors two weeks after the last daily session **(Fig S1A)**, VNS showed fewer rejected trials compared with visit 1 and visit F in CON (group x visit: F_10, 42_ = 1.74, *p* = 0.1044; visit F vs. 1 for VNS: MD = -1.5 [-2.9, 0.0], *t*(41.29) = -3.48, *p*_adj_ = 0.0643, 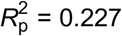; VNS vs. CON for visit F: MD = -1.3 [-2.5, -0.2], *t*(35.55) = -2.33, *p*_adj_ = 0.0257, 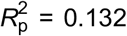 **Fig 2D**). Greater difficulty (C to N to I) increased RTs and error/lapse rates (RT: F_2, 15_ = 223.33; errors: F_2, 257_ = 29.29; lapses: F_2, 65_ = 17.10; all *p <* 0.0001; **Fig 2E**). In VNS, decreases in rejections over time were resolved to higher difficulty (I) trials (group x sex x visit x type: F_80, 28_ = 1.66, *p* = 0.0656; simple effects for men and women: each *p* < 0.0001).

**Figure 2.**
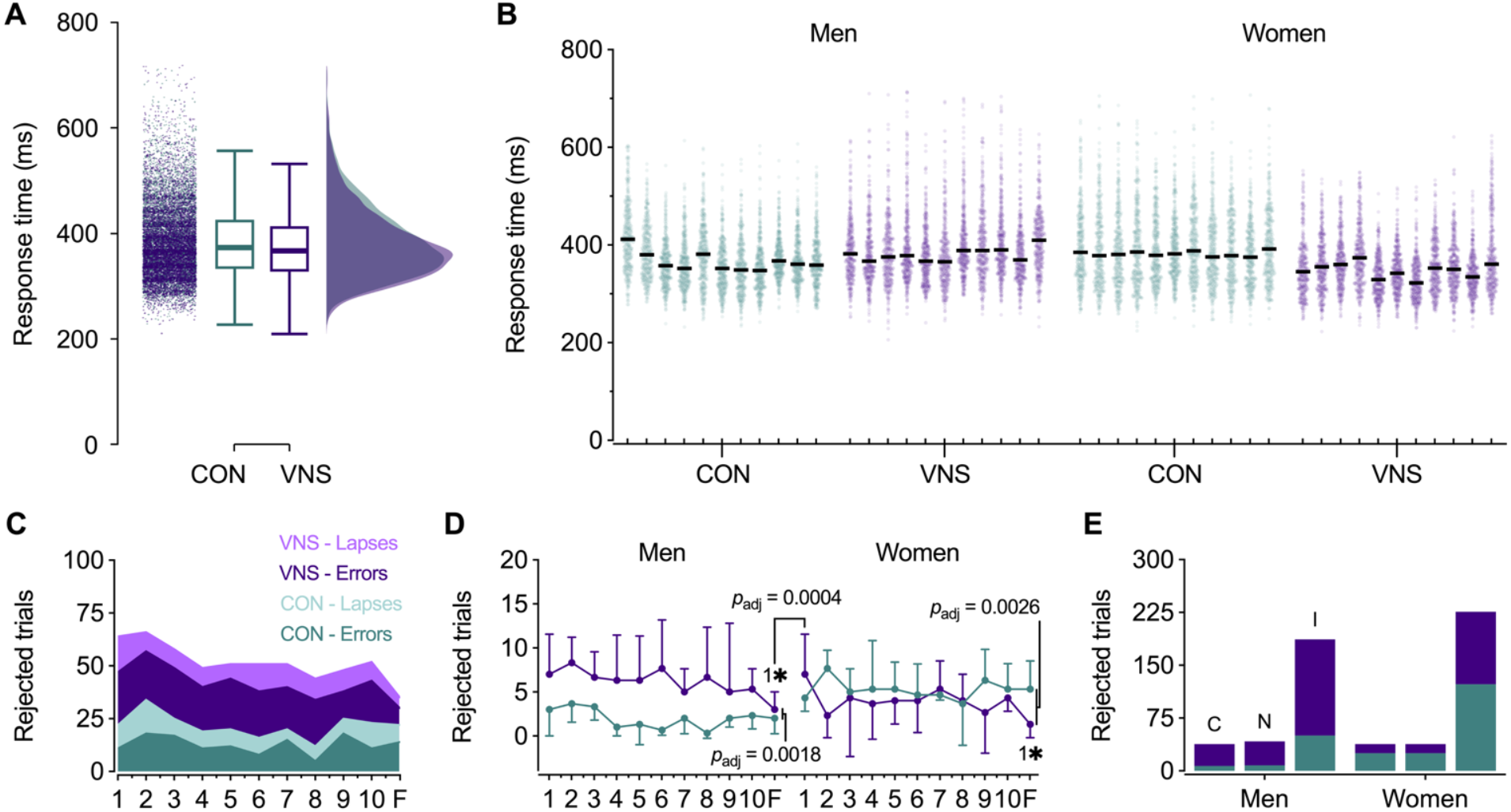
Flanker Task. **A)** Raincloud plot for Flanker response times show (*left*) retained trials (N = 15,271 after removal of 207 lapses and 362 errors); (*middle*) boxplots with medians (CON = 370 ms, VNS = 363 ms), interquartile ranges, and normative limits (1.5 x IQR); and (*right*) distributions for CON and VNS (*n* = 6 per group with 3 women each). **B)** Scatter plot with retained trials by group, sex, and visit. Data are individual trials and visit medians (black lines). **C)** Stacked area chart shows total number of errors and lapses by group and visit. **D)** Error and lapse counts (means ± SD) by group, sex, and visit. **E)** Rejected trial counts by group, sex, and stimulus type. Pairwise differences (*p*_adj_ ≤ 0.10) are indicated by Šidák-adjusted *p* values, asterisks (between-group), and numbers (within-group) after linear mixed modeling. F, follow-up visit; C, congruent; N, neutral; I, incongruent.

#### WCST

After excluding 460 lapses (2.9%) and 2,222 errors (14%, P errors = 531), 13,158 responses remained (**Fig 3A**). Median RTs decreased over time (F_10, 40_ = 2.43, *p* = 0.0229), and displayed sex-specific between-group differences (group x sex: F_1, 4_ = 6.90, *p* = 0.0597). Women in VNS responded faster than women in CON (MD = -267.6 ms [-556.0, 20.9], *t*(3.91) = -2.60, *p*_adj_ = 0.0615, 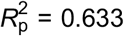); **Fig 3B**). At follow-up, RTs showed no treatment- or sex-specific differences. Errors and lapses decreased over time collectively (F_10, 165_ = 6.30, *p* < 0.0001; **Fig 3C**) but differed by group and sex (group x sex: F_1, 4_ = 14.87, *p* = 0.0181). Men in VNS had more rejected trials than men in CON (MD = 3.5 [0.6, 6.5], *t*(4.01) = 3.30, *p*_adj_ = 0.0297, 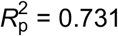 **Fig 3D**), whereas women in VNS had fewer rejections than women in CON (MD = -3.9 [-7.9, 0.1], *t*(4.01) = -2.72, *p*_adj_ = 0.0529, 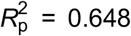), resulting in a sex-specific divergence within VNS (MD = -4.1 [-8.7, 0.5], *t*(4.01) = -2.45, *p*_adj_ = 0.0706, 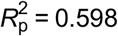). These differences reflected lower NP and P rejection rates versus women in CON (group x sex x type: F_2, 165_ = 5.51, *p* = 0.0048; NP: MD = -4.5 [-8.4, -0.6], *t*(5.25) = -2.90, *p*_adj_ = 0.0318, 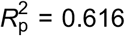 P: MD = -3.4 [-7.3, 0.5], *t*(5.25) = -2.18, *p*_adj_ = 0.0784, 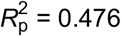 **Fig 3E**), and fewer NP rejections than men in VNS (MD = -5.7 [-10.2, -1.2], *t*(4.95) = -3.24, *p*_adj_ = 0.0234, 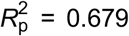). At follow-up, men in VNS exhibited further reductions in rejections (group x sex x visit: F_40, 165_ = 1.87, *p* = 0.0180; visit F vs. 1: MD = -7.8 [-14.0, - 1.7], *t*(165) = -4.28, *p*_adj_ = 0.0017, 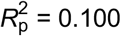 **Fig 3D**).

**Figure 3.**
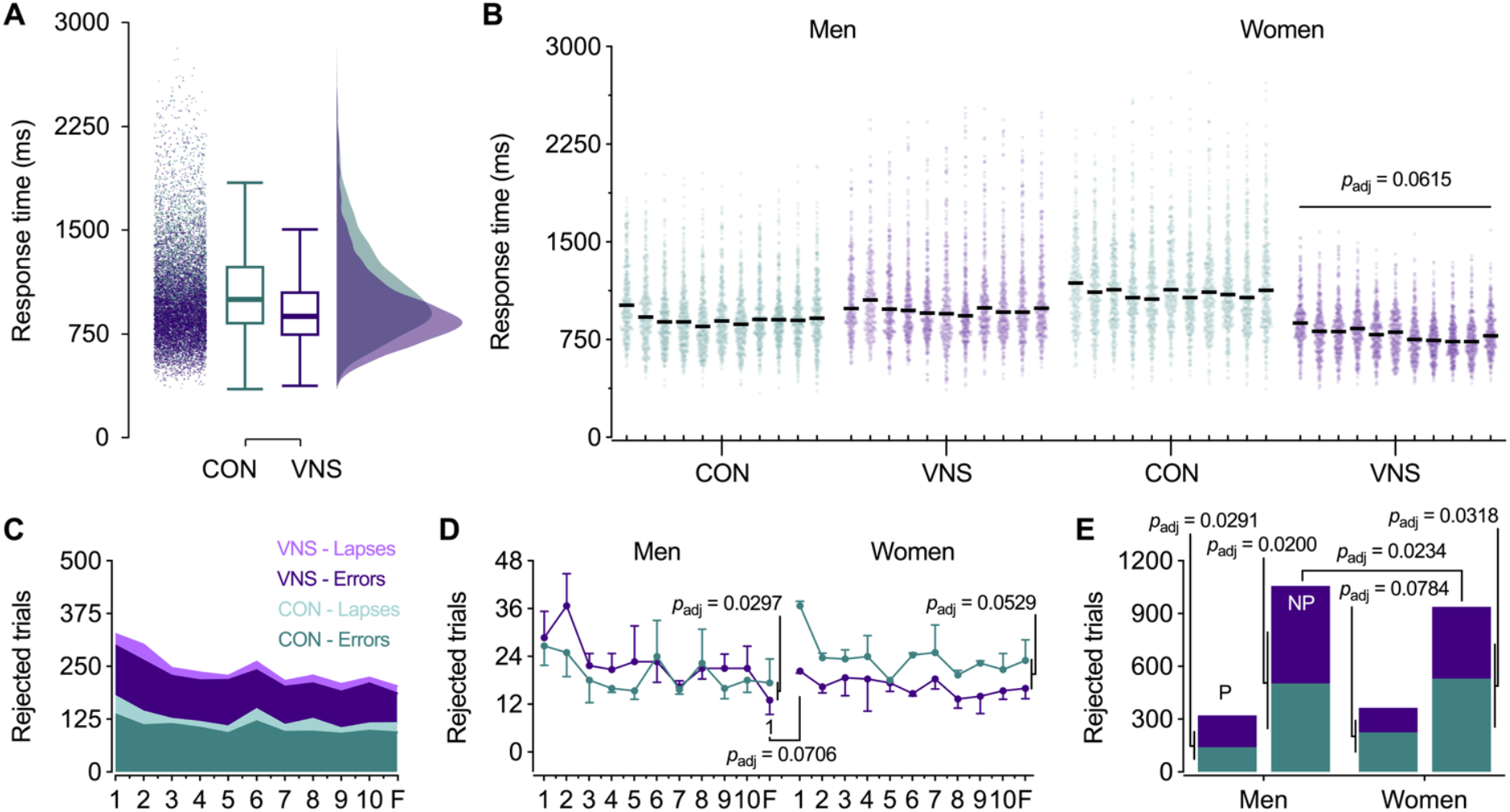
Wisconsin Card Sorting Test. **A)** Raincloud plot for WCST response times show (*left*) retained trials (N = 13,158 after removal of 460 lapses and 2,222 errors); (*middle*) boxplots with medians (CON = 989 ms, VNS = 867 ms), interquartile ranges, and normative limits (1.5 x IQR); and (*right*) distributions for CON and VNS (*n* = 6 per group with 3 women each). **B)** Scatter plot with retained trials by group, sex, and visit. Data are individual trials and visit medians (black lines). **C)** Stacked area chart shows total number of errors and lapses (rejected trials) by group and visit. **D)** Rejected trial counts (means ± SD) by group, sex, and visit. **E)** Rejected trial counts by group, sex, and error type. Pairwise differences (*p*_adj_ ≤ 0.10) are indicated by Šidák-adjusted *p* values and numbers (within-group) after linear mixed modeling. F, follow-up; P, perseverative; NP, non-perseverative.

#### SWMT

After removing 375 lapses (4.7%) and 325 errors (4.1%), 7,220 responses remained (**Fig 4A**). Median RT decreased over time (F_10, 63_ = 3.93, *p* = 0.0003), especially for women in VNS (group x sex x visit: F_20, 63_ = 1.59, *p* = 0.0834; simple effects: *p* = 0.0131; **Fig 4B**), who also displayed faster RTs than women in CON at follow-up (MD = -274.3 ms [-602.2, 53.6], *t*(5.29) = -2.12, *p*_adj_ = 0.0850,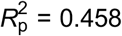). Errors and lapses decreased over time (errors: F_10, 124_ = 2.48, *p* = 0.0095; lapses: F_10, 237_ = 1.88, *p* = 0.0482; **Fig 4C**) and displayed sex-specific between-group differences (group x sex: F_1, 21_ = 20.85, *p* = 0.0002). Men in VNS had more rejected trials than men in CON (MD = 2.1 [1.3, 2.9], *t*(21.87) = 5.26, *p*_adj_ < 0.0001, 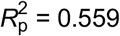 **Fig 4D**), whereas women in VNS had fewer rejections than women in CON (MD = -1.2 [-2.3, -0.1], *t*(20.42) = -2.18, *p*_adj_ = 0.0409, 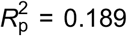) and men in VNS (MD = -2.2 [-3.4, -0.9], *t*(20.06) = -3.53, *p*_adj_ = 0.0021,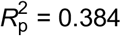).

**Figure 4.**
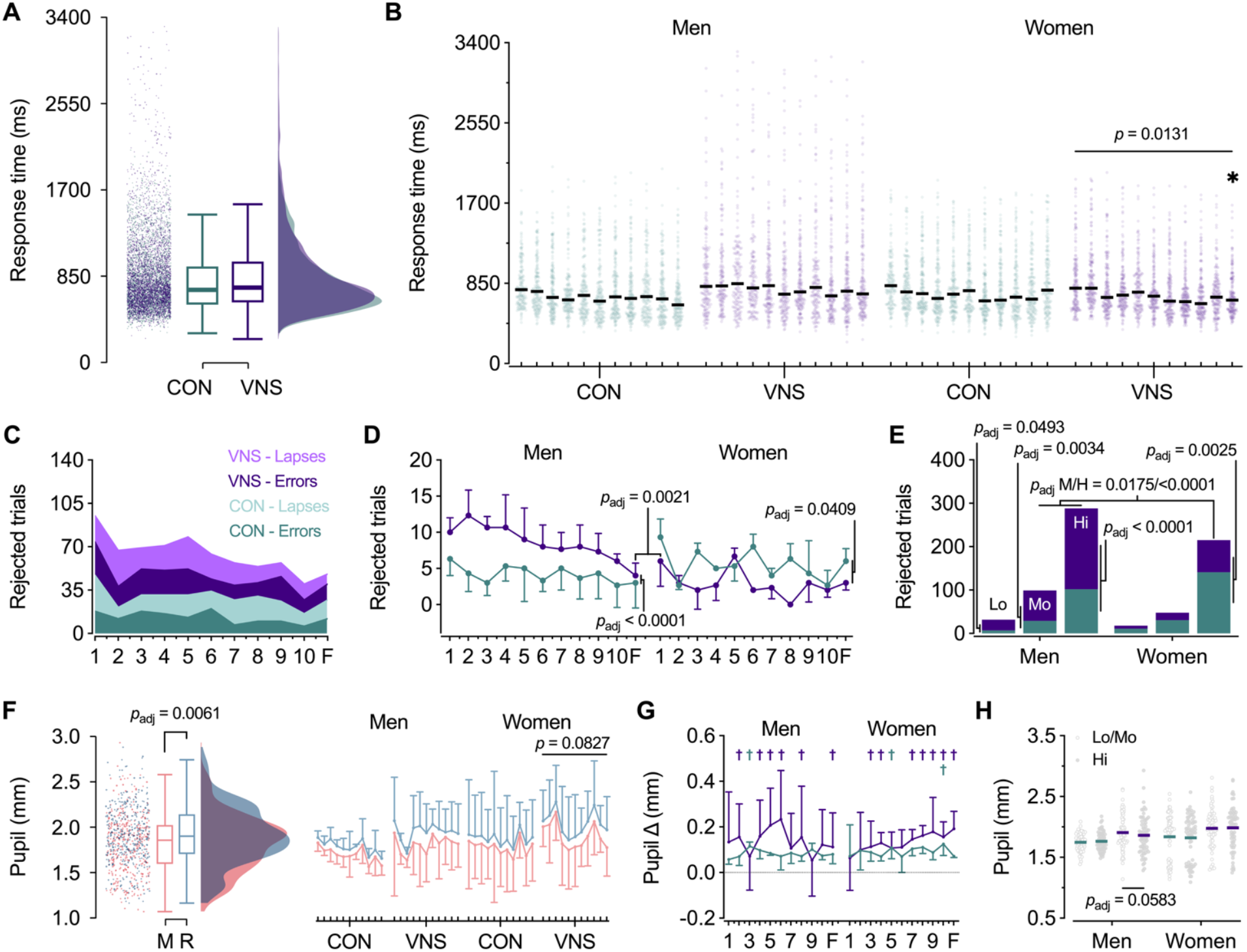
Sternberg Working Memory Task. **A)** Raincloud plot for SWMT response times show (*left*) retained trials (N = 7,220 after removal of 375 lapses and 325 errors); (*middle*) boxplots with medians (CON = 710 ms, VNS = 735 ms), interquartile ranges, and normative limits (1.5 x IQR); and (*right*) distributions for CON and VNS (*n* = 6 per group with 3 women each). **B)** Scatter plot with retained trials by group, sex, and visit. Data are individual trials and visit medians (black lines). **C)** Stacked area chart shows total number of errors and lapses (rejected trials) by group and visit. **D)** Rejected trial counts (means ± SD) by group, sex, and visit. **E)** Rejected trial counts by group, sex, and difficulty. **F)** Raincloud plot for pupil size during memorize (M) and recall (R) trial phases (*left*) along with grand means (± SD) by group, sex, phase, and visit (*right*). **G)** Task-evoked pupil response (M to R) by group, sex, and visit (MD ± SD). **H)** Pupil size by group, sex, and trial difficulty. Data are individual trials and means (black lines). Pairwise differences (*p*_adj_ ≤ 0.10) are indicated by Šidák-adjusted *p* values, asterisks (between-group), and daggers (M to R pupil size change, colored by group) after linear mixed modeling. F, follow-up; Lo, low; Mo, moderate; Hi, high.

Greater difficulty (Lo to Mo to Hi) was associated with slower RTs and higher rejection rates generally (RT: F_2, 16_ = 32.54; errors: F_2, 16_ = 40.59; lapses: F_2, 16_ = 22.58; all *p <* 0.0001). Between-group differences in rejected trials were greatest for Hi trials in men (group x sex x type: F_4, 24_ = 4.53, *p* = 0.0071; MD = 3.2 [2.0, 4.4], *t*(23.07) = 5.49, *p*_adj_ < 0.0001, 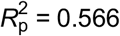 **Fig 4E**) and women (MD = -2.3 [-3.8, -0.9], *t*(21.79) = -3.42, *p*_adj_ = 0.0025, 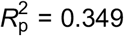). Correspondingly, decreases in rejections over time in VNS reflected improvements on Hi trials (group x sex x visit x type: F_80, 41_ = 1.45, *p* = 0.0952; simple effects for men and women: *p* < 0.0001 and 0.0064). Pupil diameter increased from memorization to recall, confirming sensitivity to cognitive load (F_1, 8_ = 13.68; MD = 0.09 mm [0.04, 0.15], *t*(8) = 3.65, *p* = 0.0061,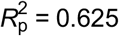 **Fig 4F)**. Pupil size did not differ by group, sex, visit, or difficulty overall (all *p* ≥ 0.5697). There was indication of greater variability among women in VNS (group x sex x visit x task: F_10, 436_ = 1.93, *p* = 0.0394; simple effects: *p* = 0.0827). In addition, compared with CON, individuals in VNS more often showed increases in pupil size from memorize to recall (59.1% vs. 13.6% of visits; **Fig 4G**). Notably, men in VNS displayed smaller pupil size during Hi trials compared with Mo and Lo trials (group x sex x type: F_2, 16_ = 3.27, *p* = 0.0647; MD = -0.05 mm [-0.09, 0.00], *t*(8) = -2.25, *p*_adj_ = 0.0583, 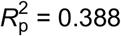 **Fig 4H**).

### Autonomic Physiological Activity

Given the absence of sex-specific autonomic responses to taVNS (pupil, SCL, mRR: tonic, *p* = 0.6788, 0.7012, 0.4410; phasic, *p* = 0.2481, 0.6226, 0.9118; **Fig S2**), aggregated results are presented. Tonic pupil size decreased across segments (F_2, 17_ = 14.45, *p* = 0.0002) but did not differ by group (*p* = 0.4794; **Fig 5A**) or group x visit (*p* = 0.6403; **Fig 5B**). Conversely, SCL was greater in VNS than CON during and after stimulation (group x segment: F_2, 16_ = 3.42, *p* = 0.0580; Stim: MD = 6.02 µS [-0.09, 12.14], *t*(4.81) = 2.56, *p*_adj_ = 0.0524, 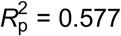 Post: MD = 7.07 µS [0.96, 13.19], *t*(4.81) = 3.01, *p*_adj_ = 0.0313, 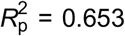 **Fig 5C/D** and **Fig S2B**), but this between-group difference did not vary across visits (*p* = 0.5514). Longer mRR was evident in VNS (group x segment: F_2, 18_ = 2.72, *p* = 0.0937; **Fig 5E**) but did not survive simple-effects resolution at Pre, Stim, or Post and did not vary across visits (*p* = 0.4634; **Fig 5F**). Respiration frequency (0.23 ± 0.02 Hz) was similar among groups (*p* = 0.7371), sexes (*p* = 0.9754), segments (*p* = 0.2904), and visits (*p* = 0.4060). Systolic and diastolic BP also showed no treatment effects **(Fig S2D**).

**Figure 5.**
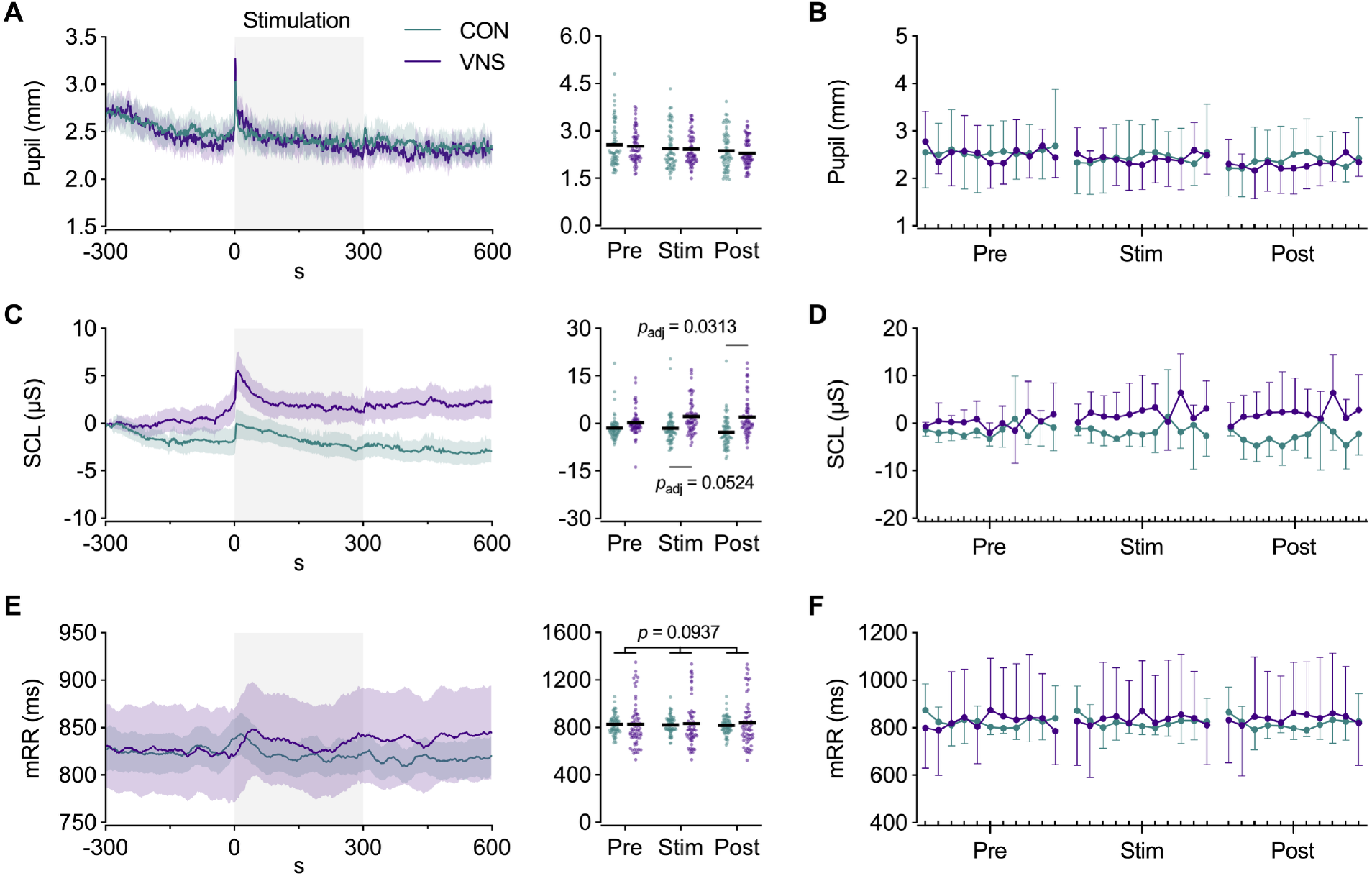
Tonic Autonomic Physiological Responses to taVNS. Pupil diameter, skin conductance level (SCL), and electrocardiographic activity (mRR intervals) were measured continuously before (Pre), during (Stim), and after (Post) five minutes of active (VNS) and sham (CON) taVNS at rest (*n* = 6 per group with 3 women each). **A)** Pupil size was similar between groups and **B)** did not vary across visits. **C)** Compared with CON, SCL was greater in VNS during and after stimulation. **D)** Increases in SCL during and after active stimulation were similar across visits. **E)** Segmental increases in mRR were evident in VNS. **F)** taVNS-induced increases in mRR were similar across visits. Data are means ± 95% CI (time series), individual segment means with grand means (black lines), and grand means ± SD. Pairwise differences (*p*_adj_ ≤ 0.10) are indicated by Šidák-adjusted *p* values after linear mixed modeling.

Stimulus-evoked pupil responses (SEPR) were greater in VNS than CON (F_1, 81_ = 20.19; MD = 0.50 mm [0.28, 0.73], *t*(81) = 4.49, *p* < 0.0001; 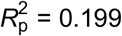 **Fig 6A**) from 0.77 to 2.92 s after stimulation onset (cluster mass = 477.1, *p*_adj_ = 0.0291), and greater in women than men overall (F_1, 81_ = 4.73; MD = 0.29 mm [0.03, 0.57], *t*(81) = 2.18, *p* = 0.0325, 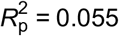) but did not differ between groups across visits (*p* = 0.1130). Time-to-peak (TTP) pupil dilation was 1.4 s (grand mean) and did not differ by group (*p* = 0.3015). Skin conductance responses (SCR) were also larger in VNS than CON (F_1, 4_ = 4.80; MD = 4.57 µS [-1.12, 10.26], *t*(4.18) = 2.19, *p* = 0.0907, 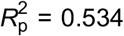 **Fig 6B**) with no difference across visits (*p* = 0.1475). SCR TTP was 6.5 s and did not differ by group (*p* = 0.1741). Stimulus-evoked mRR (SEmRR) response magnitudes did not differ by group (*p* = 0.1512) or group x visit (*p = 0*.*1942*; **Fig 6C**), but time series cluster testing identified longer SEmRR 39-146 s after stimulation onset in VNS when stimulation intensity was *not* included in the model (cluster mass = 964.5, *p*_adj_ = 0.0135). SEmRR TTP was 32 s, with no difference between groups (*p* = 0.2287).

**Figure 6.**
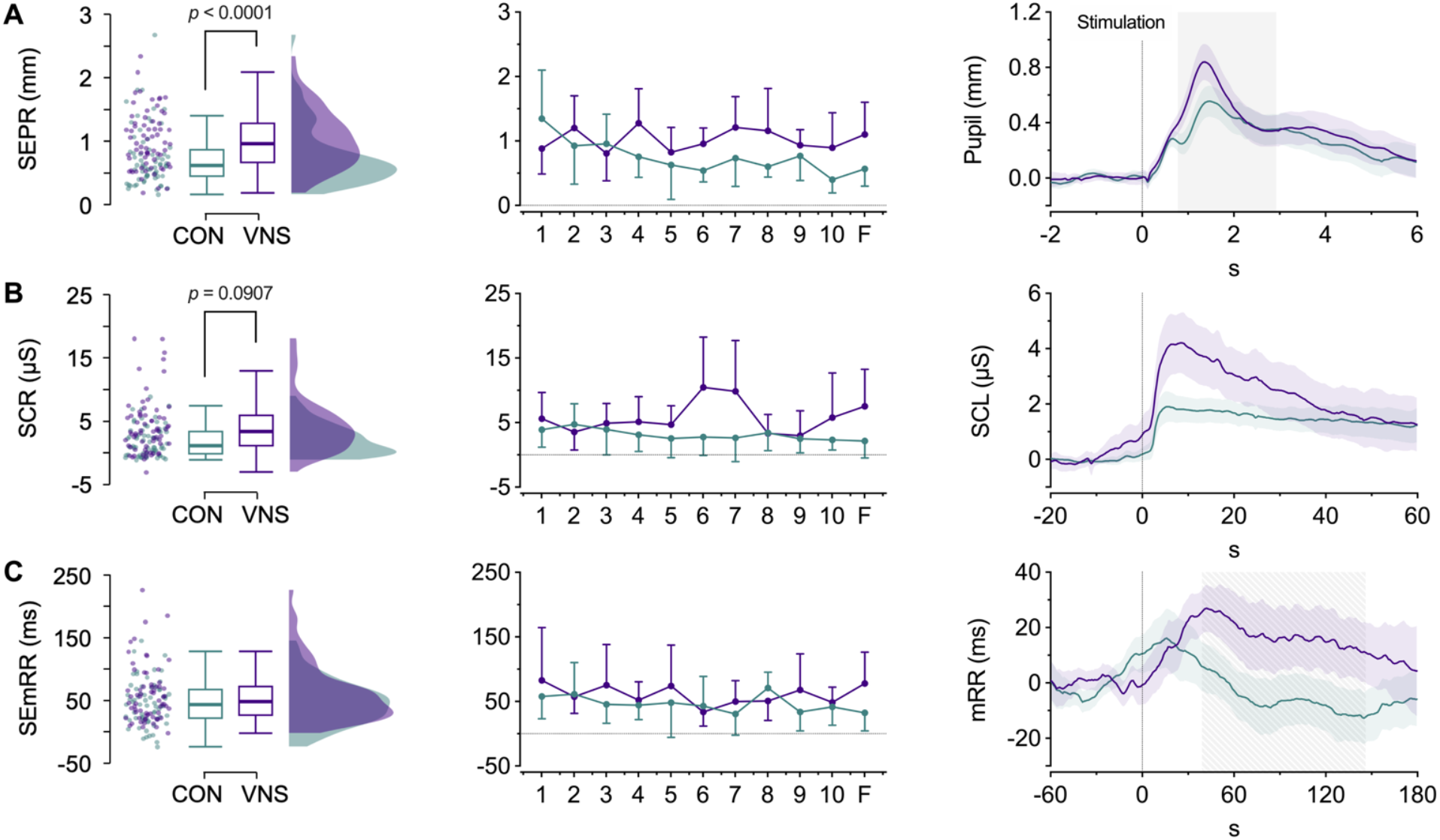
Phasic Autonomic Physiological Responses to taVNS. **A)** Stimulus-evoked pupil (SEPR), **B)** skin conductance (SCR), and **C)** mRR (SEmRR) response magnitudes by group (*left*), group x visit (*middle*), and waveforms (*right*) (*n* = 6 per group with 3 women each). SEPR magnitudes were greater in VNS than CON overall, with increases from 0.77 to 2.92 s after stimulation onset (shaded area). SCR magnitudes were also greater in VNS than CON, but differences were not resolved to a specific time-range. SEmRR magnitude did not differ between groups, but taVNS transiently increased SEmRR responses from 39-146 s in a sensitivity analysis excluding intensity covariates (hatched shaded area). Raincloud plots show (*left*) individual responses; (middle) boxplots with medians, interquartile ranges, and normative limits (1.5 x IQR); and (*right*) distributions for each group overall. Group x visit response magnitudes are means ± SD. Times series data are means ± 95% CI (N = 66 trials per group). Between-group differences (*p* ≤ 0.10) are indicated by *p* values (response magnitudes) and shaded areas (time series) for linear mixed models and non-parametric permutation tests (group x sex x visit + age (mean centered) + I*z*_*between*_ + I*z*_*within*_ + error (ID / visit)) with cluster mass-based correction, respectively.

### Discomforts, Perceived Effects, and Blinding

Stimulation was well tolerated: no sessions were terminated early, no dose reductions were required, and pain induction was not reported. All adverse events were minor and self-limited, with no evidence of sex specificity. Across 132 encounters, headache was reported on five occasions (3.8%). The only other reported discomfort was minor skin irritation at electrode sites (*n* = 17 instances; 12.9% of visits). Perceived effects varied but were more likely for active taVNS: 5/6 VNS participants reported treatment effects on some (*n* = 5) or all (*n* = 1) visits, whereas 5/6 CON participants reported no treatment effects on some (*n* = 2) or all (*n* = 3) visits. Blinding was reasonably successful: in VNS, 3/6 could not identify condition, while 3/6 always (*n* = 2) or mostly (*n* = 1) correctly guessed active; in CON, 4/6 could not identify condition, 1/6 correctly identified sham, and 1/6 consistently guessed active. Bang’s Blinding Index was 0.52 [0.39, 0.64] for VNS and 0.06 [-0.11, 0.24] for CON [55].

## DISCUSSION

We asked whether ten consecutive days of monoaural taVNS can promote immediate and durable cumulative changes in executive function and biomarkers of proposed neuromodulatory mechanisms in healthy young adults. In addition to replicating growing evidence that executive function and performance accuracy are responsive to acute taVNS, our results indicate that daily taVNS can promote load-dependent cognitive effects that accumulate over time and, in several instances, persist at a two-week rechallenge. Stimulation-induced changes in resting pupil size, skin conductance, and cardiorespiratory activity were evident in aggregate but variable, and we did not detect group- or sex-specific longitudinal changes in these autonomic measures. Nevertheless, the convergence of behavioral and autonomic responses to taVNS is consistent with the proposed modulation of adaptive gain in task-relevant networks through stimulation-induced changes in LC-NE, BFCN, or GABAergic signaling [11, 13, 31, 86, 87]. This study addresses three gaps in the evidence base: a repeated-dosing approach with dense high-fidelity sampling, incorporation of distinct executive function subdomains, and explicit consideration of sex as a biological variable.

### Cognitive Function

The vast majority of taVNS studies have emphasized single sessions and produced inconsistent cognitive effects. Our observations suggest that dense repeated sampling conferred two advantages. First, daily assessments enabled the emergence, stabilization, and accumulation of effects that were absent or transient on the first visit. Second, this approach reduced measurement noise and within-person variability, thereby increasing sensitivity, and revealing interactions with load and sex that would otherwise be obscured. In accordance with recent meta-analysis [25], the most pronounced cognitive effects of longitudinal taVNS were resolved to performance accuracy. Women in VNS displayed fewer Flanker trial rejections overall, with further improvements in accuracy at follow-up. Among men and women in VNS, slower Flanker response times at follow-up coincided with improvements in accuracy, consistent with a speed-accuracy trade-off, rather than an adverse residual treatment effect. On the WCST, women in VNS displayed immediate and sustained decreases in response times and rejections, including perseverative errors, extending evidence that acute taVNS can enhance shifting performance [28]. Women in VNS also showed sustained gains on the SWMT, where benefits emerged as fewer high-difficulty rejections and shorter response times. Across tasks, men who received active taVNS experienced load-dependent increases in errors and lapses and no comparative differences or changes in response speed. However, men in VNS also displayed larger decreases in rejections over time and similar behavioral performance at follow-up. Our results thus provide preliminary evidence to support the dissociation of acute and longitudinal taVNS cognitive effects.

Mechanistically, cognitive responses to taVNS are consistent with the premise that it engages overlapping FPN and SN nodes that play central roles in inhibition (Flanker), shifting (WCST), and WM (SWMT) [59-64, 88, 89]. Neuroimaging studies have repeatedly implicated these networks in taVNS effects based on stimulation-induced changes in functional activity and connectivity [7, 9, 10, 12, 90, 91]. In addition to the lower likelihood of behavioral responses to a single session of stimulation, the inconsistency of acute taVNS cognitive effects likely reflects variation in interactions between stimulation-induced neurophysiological activity and task-related activity specific to different cognitive domains, subdomains, and task-variants [11, 13, 19, 74]. For example, and similar to other executive function subdomains [60, 62, 89], WM task-variants are associated with overlapping *and* distinct functional activity patterns [59, 61, 63, 92]. Accordingly, acute taVNS improved performance on spatial-but not digit-based n-back working memory tests [21]; both of which emphasize distinct WM sub-mechanisms compared with each other (visuospatial vs. verbal) and the SWMT (manipulation vs. maintenance) [93]. Taken together, the specificity of taVNS cognitive effects represents a unique challenge and opportunity for efforts to understand and improve this non-invasive brain stimulation technique.

Regardless of executive function subdomain, our results support growing evidence that cognitive responses to taVNS increase with task difficulty [21, 22, 28, 29]. When cognitive load increases with inhibitory requirements (Flanker incongruent trials), overall task difficulty (WCST), or WM load (SWMT Hi trials), performance depends on the dynamic allocation of cognitive resources, stabilization of task-relevant representations, and suppression of distractions. Load-specific taVNS effects may reflect the greater dependence of high-load trials and tasks on adaptive gain [86], with taVNS plausibly increasing gain through its neuromodulatory effects on task-relevant FPN and SN pathways when cognitive load and stimulation intensity fall within an optimal arousal range [11, 13, 74, 91]. On the Flanker, the resolution of longitudinal improvements in accuracy to incongruent trials is consistent with the facilitation of inhibition mediated by the right vlPFC, which displays increased functional activity in response to acute taVNS [7, 9, 94]. In accordance with slower response times and lower accuracy compared with the Flanker and SWMT, the WCST is a relatively complex task that involves contributions from numerous executive function subdomains [62, 64, 88, 89]. Comparatively stronger taVNS effects on WCST performance may thus reflect enhanced neuromodulatory input to relevant fronto-striatal-thalamic loops during sustained feedback-guided updating [11, 29, 86-88]. On the SWMT, in addition to the resolution of taVNS cognitive effects to higher difficulty trials, stimulation-induced increases in task-evoked pupil responses are compatible with the enhancement of neuromodulatory input to FPN and SN nodes that facilitate load-dependent maintenance and retrieval processes [59, 61-63, 93]. The persistence of enhanced behavioral and pupil responses at follow-up raises the possibility that repeated taVNS induces durable synaptic or circuit-level adaptations in these systems and their targets.

While underpowered for definitive conclusions, the results of this investigation add to growing evidence that sex can be an important factor for neuromodulation effects [36, 42, 43, 95]. Behavioral and autonomic outcomes were similar during the first visit and at baseline, which suggests that sexually dimorphic responses to daily taVNS were not the result of differences in homeostatic or ceiling effects. Given that women had lower perceptual sensory thresholds, and performed better when intensity was comparatively lower, the pattern of effects was compatible with sex-dependent dose sensitivity (see **S Results**). In addition, studies with invasive VNS indicate that stimulation efficacy follows an inverted U-shaped (Yerkes-Dodson) function, with moderate intensities producing the largest effects [96, 97]. Thus, one potential explanation for the observed cognitive responses to taVNS is that differences in absolute intensity placed men and women at distinct points on the Yerkes-Dodson/Hardy-Fazey curve, with women positioned closer to optimal arousal and men toward over-arousal [98-100]. Accordingly, men in VNS displayed worse accuracy and smaller pupil size on high-load SWMT trials *specifically*. Greater intensities could also induce differential effects secondary to the recruitment of higher threshold (i.e., Aδ or C fibers) or non-ABVN cutaneous afferents [8, 101]. The stimulation parameters used in this study were designed to avoid pain induction, and participant reports support the achievement of this goal [5, 102, 103]. It is not clear whether sex-specific differences in sensory thresholds are an idiosyncratic feature of this sample or generalizable [45, 90, 104]. Alternatively and in addition to endogenous or contraceptive-related neurosteroid concentrations [37-39], sex-specific cognitive effects could reflect differences in the structural properties of frontoparietal areas that may mediate taVNS effects on executive function [40, 41]. The neurophysiological basis of sexual dimorphism requires clarification, but our results provide further reason to consider sex as a biological variable in VNS research.

### Autonomic Physiological Activity

Tonic pupil size steadily decreased from the beginning to end of each resting assessment and was unaffected by 10 consecutive days of taVNS, confirming and extending evidence that acute taVNS does not increase tonic pupil size at rest [32, 33, 105, 106]. In contrast, SCL increased during and after stimulation, consistent with activation of the NTS-paraventricular nucleus (PVN) pathway and downstream excitation of spinal intermediolateral sympathetic preganglionic neurons via direct projections from the PVN or parallel pathways involving the rostral ventrolateral medulla (RVLM) [18]. Stimulus-evoked pupil and skin conductance responses were larger among individuals in VNS, extending evidence that acute taVNS increases LC-NE-mediated autonomic activity [33, 71, 73, 105, 107]. At the same time, taVNS produced a time-localized lengthening of mRR that disappeared when stimulation intensity was included as a covariate, suggesting that bradycardic responses depend on dose and baseline cardiac state, and providing another plausible explanation for the heterogeneity of cardiorespiratory effects to date [3, 31, 32, 34, 42, 44, 45, 108-112]. In the absence of direct evidence for changes in cardiovagal drive, NTS projections to GABAergic caudal ventrolateral medullary neurons that inhibit RVLM cardiac sympathetic outflow offer a plausible route for transient mRR lengthening [3, 18, 34, 42, 113]. Given the sensitivity of sympathetic and cardiovagal outputs to respiratory activity [18], the direct confirmation of equivalence across groups, sexes, segments, and visits supports the conclusion that differences in autonomic activity were not driven by respiratory confounding.

Collectively, the observed autonomic physiological responses to daily taVNS are consistent with the enhancement of autonomic responsiveness, predominantly via phasic effects (SEPR, SCR, transient mRR lengthening), alongside a modest sustained increase in arousal (SCL) during and after stimulation. These dual effects coincide with responses to invasive VNS, but because taVNS exclusively targets afferent fibers, peripheral changes are necessarily indirect and state-dependent [8, 18, 31]. In addition, the finding that stimulus-evoked pupil and skin conductance responses scale with stimulation intensity and sex, while tonic pupil size remains stable, reinforces the ideas that 1) resting LC-NE activation is brief and dose-dependent; 2) longer-lasting effects likely involve additional neuromodulatory systems such as BFCN and NTS-GABA; and 3) failure to account for differences in stimulation dose and sex may confound or suppress treatment effects. In animal models, noradrenergic lesions abolish SCRs [114], underscoring a noradrenergic contribution to sudomotor responses; translation to humans should be made cautiously. Nevertheless, in the context of taVNS effects on executive function, it is notable that prominent neural correlates of SCL include various CAN, SN, and FPN nodes [115]. The hybrid event-related block design used to assess resting autonomic responses to taVNS allowed for concurrent assessment of slow, continuous changes and transient stimulus-evoked responses, eliminating a major source of non-systematic variability.

### Limitations and Future Directions

While guided by *a priori* hypotheses, considering the small sample and sex-balanced but underpowered subgroup tests, our findings are exploratory. Task order was fixed, blinding was single, and stimulation was monoaural. The earlobe sham is widely used but not physiologically inert and can elicit modest central and autonomic activation [7-9, 28, 32, 101, 116]. Greater absolute current in sham did not reproduce the pattern seen with active taVNS, and inclusion of intensity as a covariate often strengthened active-sham differences, which suggests that the observed taVNS effects are not attributable to confounding from sensory-driven orienting or startle responses. Given a smaller number of trials and greater variability, determination of autonomic adaptive responses to daily taVNS will require a larger sample or different experimental approach. Finally, the study was not designed to adjudicate among neuromodulatory or cognitive mechanisms: inferences rest on converging but indirect evidence and precedence. In addition to replication in a larger cohort, incorporation of imaging or neurochemical biomarkers, and systematic comparison of monoaural versus binaural stimulation, future work should precisely control or model absolute current density and interactions between sex and dose.

### Conclusions

Ten days of monoaural taVNS produced immediate and cumulative changes in executive function that remained evident at a two-week rechallenge. Cognitive effects extended to several executive function subdomains, including inhibition, shifting, and working memory. Overall, taVNS had the strongest effect on accuracy. Behavioral benefits were concentrated in women, whereas men displayed initial performance costs, but with greater longitudinal improvements in accuracy. The resolution of taVNS cognitive effects to higher difficulty trials and tasks is consistent with the modulation of adaptive gain in executive function networks. In addition, concurrent increases in stimulus-evoked pupil and skin conductance responses, transient mRR lengthening, and resting skin conductance levels suggest that acute taVNS enhanced autonomic responsiveness and arousal. The results are consistent with a model in which daily taVNS induces nonlinear and state-dependent neuromodulatory effects to facilitate adaptations that outlast stimulation duration. Our findings identify biological sex and stimulation intensity as potentially important considerations for taVNS efficacy, and provide high-fidelity longitudinal data to support dosing, biomarker selection, and trial design in future studies. Collectively, the results support continued development of taVNS as tool for the modulation of executive function and autonomic activity.

## Supporting information

Supplemental Results

## ACKNOWLEDGEMENTS

We thank the volunteers for their commitment. The opinions or assertions in this manuscript are the private views of the authors and should not be construed as official policy or reflecting the views of the United States Air Force or DEVCOM Army Research Laboratory. Any citations of trade names in this report do not constitute an official DoD endorsement or approval of products. The investigators have adhered to the policies for protection of human subjects as prescribed in DoDI3216.02_DAFI40-402, and the research was conducted in adherence with the provisions of 45 CFR Part 46.

