## Supplemental Materials for "Cognitive and autonomic physiological responses to daily transcutaneous auricular vagus nerve stimulation in healthy adults"

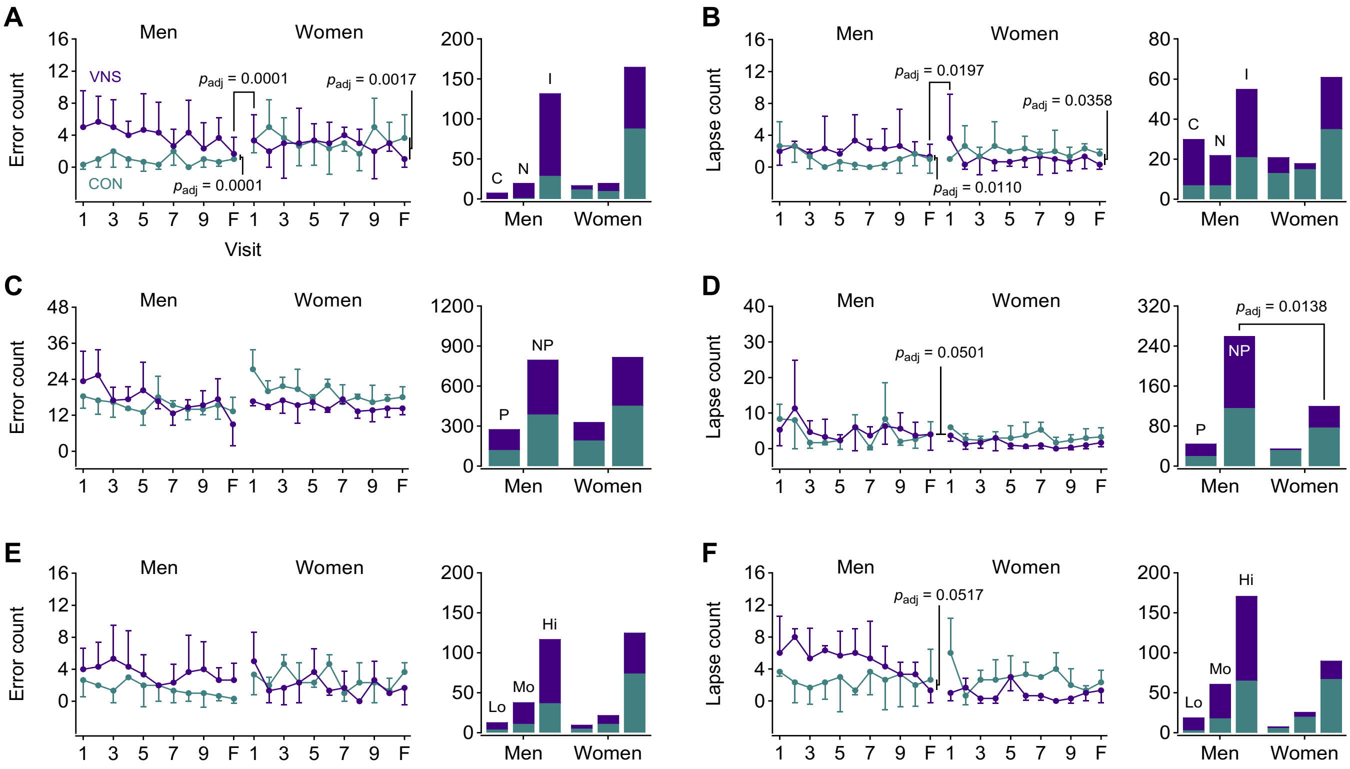


**S Figure 1. Cognitive Test Errors and Lapses.** Error and lapse counts by group, sex, and visit (*left*: means ± SD), or group, sex, and type (*right*: total counts) for the Flanker (**A/B**), WCST (**C/D**), and SWMT (**E/F**). Pairwise differences ($p_{adj}$ ≤ 0.10) are indicated by Šidák-adjusted *p* values after linear mixed modeling. CON (sham); VNS (active); C, congruent; N, neutral; **I**, incongruent; P, perseverative; NP, non-perseverative; Lo, low; Mo, moderate; Hi, high.

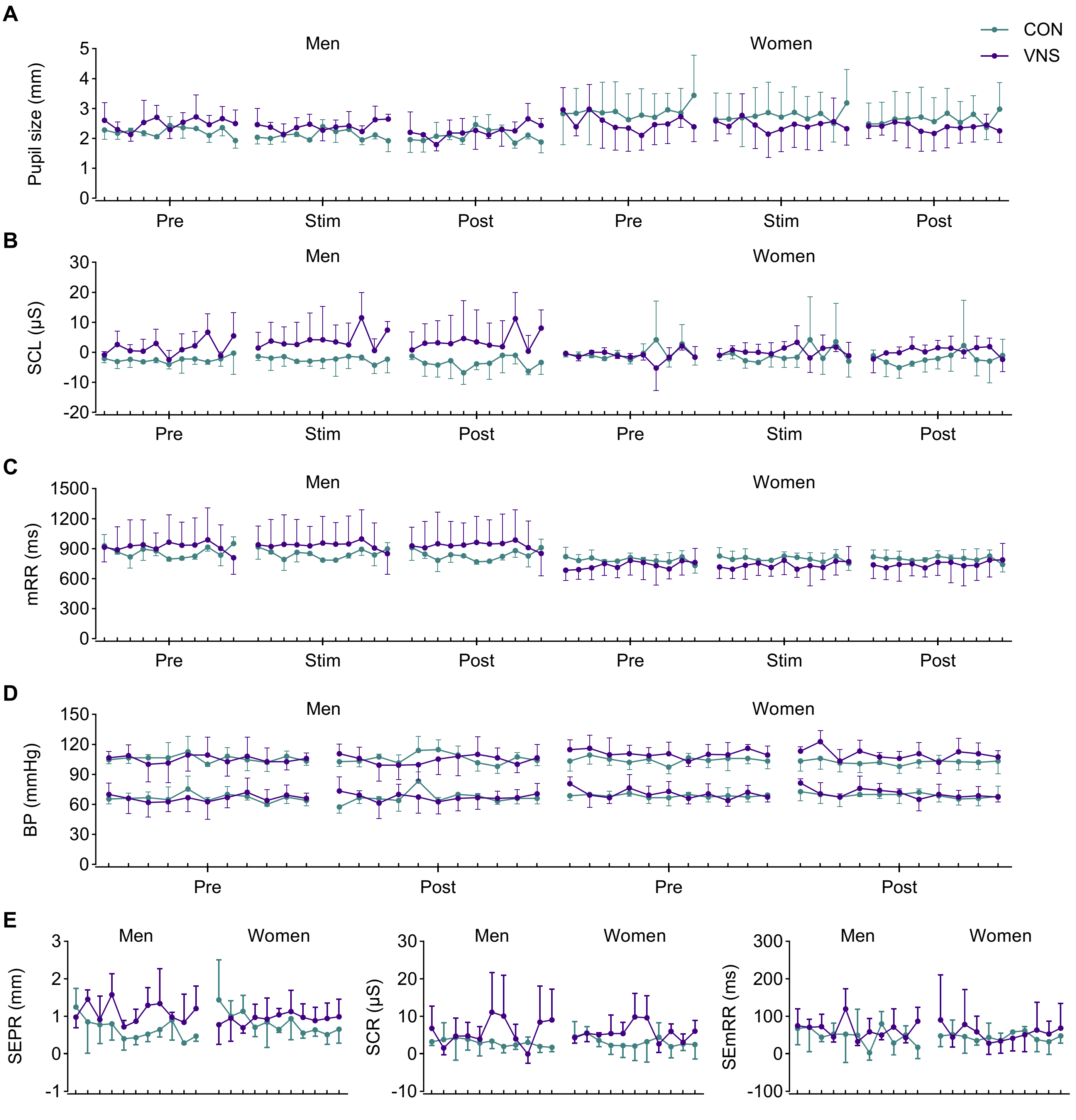


**S Figure 2. Resting Autonomic Physiological Activity.** Summarized data for each measure by group, sex, segment, and visit. CON (sham); VNS (active); SCL, skin conductance level; mRR, mean R-R interval; BP, blood pressure; SEPR, stimulus-evoked pupil response; SCR, skin conductance response; SEmRR, stimulus-evoked mRR interval.

**Supplementary Table 1. Experimental Controls**

| Visit | Group | Hydration (USG) | | | Sleep (hr) | | | Sleep quality (0-5) | | | Soreness (0-100) | | | Life stress (0-946) | | |
|---|---|---|---|---|---|---|---|---|---|---|---|---|---|---|---|---|
| 1 | CON | 1.014 | ± | 0.006 | 7.7 | ± | 1.5 | 4.0 | ± | 1.1 | 0.8 | ± | 1.2 | 61.0 | ± | 80.4 |
| | VNS | 1.029 | ± | 0.036 | 7.3 | ± | 1.5 | 3.5 | ± | 0.5 | 0.7 | ± | 1.2 | 40.8 | ± | 61.1 |
| 2 | CON | 1.016 | ± | 0.008 | 7.5 | ± | 1.1 | 3.8 | ± | 0.8 | 0.3 | ± | 0.5 | 37.2 | ± | 56.2 |
| | VNS | 1.014 | ± | 0.010 | 6.8 | ± | 0.8 | 3.2 | ± | 0.8 | 1.3 | ± | 2.8 | 32.0 | ± | 47.7 |
| 3 | CON | 1.017 | ± | 0.004 | 6.4 | ± | 1.4 | 3.3 | ± | 1.0 | 0.2 | ± | 0.4 | 43.0 | ± | 60.5 |
| | VNS | 1.013 | ± | 0.010 | 6.9 | ± | 0.9 | 3.5 | ± | 0.8 | 0.5 | ± | 0.8 | 32.5 | ± | 56.6 |
| 4 | CON | 1.016 | ± | 0.007 | 6.8 | ± | 1.2 | 3.3 | ± | 1.0 | 0.3 | ± | 0.5 | 36.3 | ± | 45.4 |
| | VNS | 1.009 | ± | 0.005 | 6.3 | ± | 1.0 | 2.8 | ± | 1.0 | 0.7 | ± | 1.0 | 33.2 | ± | 56.7 |
| 5 | CON | 1.011 | ± | 0.010 | 8.0 | ± | 0.6 | 3.7 | ± | 0.5 | 1.2 | ± | 1.2 | 40.8 | ± | 47.5 |
| | VNS | 1.010 | ± | 0.010 | 7.5 | ± | 1.6 | 3.2 | ± | 0.8 | 0.5 | ± | 0.8 | 32.0 | ± | 47.6 |
| 6 | CON | 1.020 | ± | 0.005 | 6.9 | ± | 1.0 | 3.8 | ± | 0.8 | 0.8 | ± | 1.2 | 44.3 | ± | 51.5 |
| | VNS | 1.010 | ± | 0.008 | 7.1 | ± | 1.6 | 3.5 | ± | 0.8 | 0.3 | ± | 0.8 | 36.8 | ± | 55.0 |
| 7 | CON | 1.021 | ± | 0.002 | 7.5 | ± | 1.1 | 3.8 | ± | 0.8 | 0.8 | ± | 1.2 | 47.7 | ± | 50.7 |
| | VNS | 1.013 | ± | 0.009 | 7.0 | ± | 1.2 | 3.7 | ± | 0.8 | 0.8 | ± | 0.8 | 36.8 | ± | 58.1 |
| 8 | CON | 1.019 | ± | 0.005 | 7.6 | ± | 1.1 | 3.7 | ± | 1.0 | 0.8 | ± | 1.6 | 47.7 | ± | 50.7 |
| | VNS | 1.011 | ± | 0.008 | 6.9 | ± | 1.0 | 3.3 | ± | 0.5 | 0.7 | ± | 1.2 | 32.5 | ± | 56.6 |
| 9 | CON | 1.016 | ± | 0.008 | 6.8 | ± | 0.8 | 3.7 | ± | 0.8 | 1.0 | ± | 2.0 | 47.7 | ± | 50.7 |
| | VNS | 1.015 | ± | 0.008 | 6.6 | ± | 1.9 | 3.2 | ± | 0.8 | 0.8 | ± | 1.2 | 36.8 | ± | 58.1 |
| 10 | CON | 1.014 | ± | 0.010 | 7.2 | ± | 0.8 | 3.7 | ± | 1.0 | 0.7 | ± | 0.8 | 43.5 | ± | 50.2 |
| | VNS | 1.013 | ± | 0.010 | 8.3 | ± | 2.1 | 3.8 | ± | 0.8 | 0.7 | ± | 1.2 | 36.8 | ± | 58.1 |
| 11 | CON | 1.021 | ± | 0.003 | 7.3 | ± | 0.9 | 3.8 | ± | 0.9 | 0.8 | ± | 0.9 | 28.2 | ± | 63.3 |
| | VNS | 1.015 | ± | 0.010 | 7.3 | ± | 1.1 | 3.5 | ± | 0.5 | 1.8 | ± | 1.8 | 33.8 | ± | 56.8 |
| | **Group (*p*)** | **0.3352** | | | **0.7332** | | | **0.3739** | | | **0.8628** | | | **0.7931** | | |
| | **Visit (*p*)** | **0.3561** | | | **0.1411** | | | **0.2185** | | | **0.4671** | | | **0.1391** | | |
| | **Group x Visit (*p*)** | **0.2673** | | | **0.4670** | | | **0.6193** | | | **0.4203** | | | **0.5434** | | |

Values are means ± SD. Two-way ANOVAs confirmed similar hydration, sleep quantity and quality, soreness/pain, and life stress between groups and across visits.

**Supplementary Table 2. Cognitive Test Completion Times**

| Visit | Group | Sex | Flanker | | | WSCT | | | SWMT | | |
|---|---|---|---|---|---|---|---|---|---|---|---|
| 1 | CON | Men | 799.7 | ± | 66.3 | 804.0 | ± | 66.8 | 813.0 | ± | 66.7 |
| | | Women | 585.4 | ± | 140.1 | 589.7 | ± | 140.0 | 598.7 | ± | 139.4 |
| | VNS | Men | 722.6 | ± | 197.5 | 726.7 | ± | 197.1 | 736.3 | ± | 198.8 |
| | | Women | 680.6 | ± | 254.3 | 685.3 | ± | 253.9 | 693.6 | ± | 253.6 |
| 2 | CON | Men | 707.5 | ± | 59.9 | 712.1 | ± | 59.9 | 720.9 | ± | 59.9 |
| | | Women | 590.2 | ± | 128.9 | 595.2 | ± | 128.4 | 603.7 | ± | 128.1 |
| | VNS | Men | 687.1 | ± | 174.3 | 691.0 | ± | 174.2 | 699.9 | ± | 174.0 |
| | | Women | 634.8 | ± | 133.4 | 638.9 | ± | 133.5 | 647.0 | ± | 133.6 |
| 3 | CON | Men | 782.7 | ± | 119.5 | 787.0 | ± | 119.5 | 795.3 | ± | 119.8 |
| | | Women | 530.9 | ± | 174.5 | 534.8 | ± | 174.3 | 544.3 | ± | 174.0 |
| | VNS | Men | 680.5 | ± | 168.1 | 684.2 | ± | 167.9 | 692.9 | ± | 167.8 |
| | | Women | 655.4 | ± | 165.1 | 659.6 | ± | 165.1 | 667.6 | ± | 165.2 |
| 4 | CON | Men | 739.2 | ± | 127.1 | 743.6 | ± | 127.2 | 751.6 | ± | 127.0 |
| | | Women | 535.1 | ± | 178.7 | 539.3 | ± | 178.5 | 547.9 | ± | 178.5 |
| | VNS | Men | 684.3 | ± | 185.4 | 688.5 | ± | 185.3 | 697.0 | ± | 185.3 |
| | | Women | 647.9 | ± | 165.2 | 652.1 | ± | 165.0 | 659.9 | ± | 164.7 |
| 5 | CON | Men | 758.9 | ± | 111.1 | 763.1 | ± | 111.5 | 771.2 | ± | 111.4 |
| | | Women | 571.2 | ± | 139.3 | 575.3 | ± | 139.1 | 583.8 | ± | 139.1 |
| | VNS | Men | 684.9 | ± | 175.1 | 690.6 | ± | 173.3 | 697.3 | ± | 170.2 |
| | | Women | 644.9 | ± | 159.8 | 648.7 | ± | 159.8 | 656.5 | ± | 159.4 |
| 6 | CON | Men | 779.1 | ± | 150.0 | 783.3 | ± | 150.5 | 791.6 | ± | 150.5 |
| | | Women | 573.4 | ± | 139.7 | 577.5 | ± | 139.5 | 586.1 | ± | 139.8 |
| | VNS | Men | 678.2 | ± | 176.5 | 682.2 | ± | 176.4 | 693.9 | ± | 181.4 |
| | | Women | 645.7 | ± | 160.3 | 649.7 | ± | 160.5 | 657.4 | ± | 160.1 |
| 7 | CON | Men | 759.9 | ± | 110.5 | 764.0 | ± | 111.1 | 772.0 | ± | 111.0 |
| | | Women | 570.2 | ± | 142.6 | 574.3 | ± | 142.5 | 582.8 | ± | 142.3 |
| | VNS | Men | 689.1 | ± | 175.4 | 693.1 | ± | 175.3 | 699.3 | ± | 168.6 |
| | | Women | 651.1 | ± | 167.6 | 654.9 | ± | 167.6 | 662.8 | ± | 167.6 |
| 8 | CON | Men | 758.3 | ± | 113.0 | 762.5 | ± | 113.5 | 771.5 | ± | 112.9 |
| | | Women | 576.2 | ± | 140.2 | 580.7 | ± | 140.8 | 589.3 | ± | 140.7 |
| | VNS | Men | 684.7 | ± | 169.8 | 688.6 | ± | 169.8 | 697.5 | ± | 169.7 |
| | | Women | 683.0 | ± | 223.7 | 687.0 | ± | 223.3 | 694.7 | ± | 223.0 |
| 9 | CON | Men | 753.8 | ± | 114.3 | 758.2 | ± | 114.7 | 766.4 | ± | 114.6 |
| | | Women | 515.0 | ± | 43.4 | 519.7 | ± | 41.8 | 528.4 | ± | 41.9 |
| | VNS | Men | 662.3 | ± | 191.3 | 666.3 | ± | 191.3 | 675.0 | ± | 191.3 |
| | | Women | 643.9 | ± | 164.0 | 648.4 | ± | 163.9 | 655.9 | ± | 163.8 |
| 10 | CON | Men | 688.6 | ± | 57.5 | 692.4 | ± | 57.7 | 700.9 | ± | 57.5 |
| | | Women | 579.1 | ± | 143.6 | 583.6 | ± | 144.0 | 592.3 | ± | 143.8 |
| | VNS | Men | 698.4 | ± | 157.4 | 702.2 | ± | 157.2 | 710.9 | ± | 157.3 |
| | | Women | 644.3 | ± | 161.7 | 649.0 | ± | 161.0 | 656.6 | ± | 161.0 |
| F | CON | Men | 774.0 | ± | 94.4 | 778.0 | ± | 94.7 | 786.4 | ± | 94.8 |
| | | Women | 615.1 | ± | 75.5 | 619.2 | ± | 75.3 | 628.0 | ± | 75.1 |
| | VNS | Men | 702.5 | ± | 163.0 | 706.9 | ± | 163.4 | 715.4 | ± | 163.1 |
| | | Women | 660.7 | ± | 147.9 | 664.9 | ± | 147.4 | 672.7 | ± | 147.2 |
| | | **Group (*p*)** | | **0.9074** | | | **0.9079** | | | **0.9110** | |
| | | **Sex (*p*)** | | **0.2249** | | | **0.2248** | | | **0.2234** | |
| | | **Visit (*p*)** | | **0.1684** | | | **0.1707** | | | **0.1655** | |
| | | **Group x Sex x Visit (*p*)** | | **0.3008** | | | **0.2986** | | | **0.3071** | |

Values are means ± SD (in seconds). Three-way ANOVA confirmed similar completion times among groups, sexes, and visits.

### Supplementary Table 3. Linear Mixed Model Optimization

#### A. Flanker Task

| Model | Data | Method | Fixed (FE) | Random (Cov) | Repeated (Cov) | Param | BIC | AIC | -2 LL |
|---|---|---|---|---|---|---|---|---|---|
| 1 | medRT | ML | Age Group Sex Visit Type Group*Sex Group*Visit Group*Sex*Visit Group*Type Group*Sex*Type Group*Sex*Type*Visit | None | None | 134 | 4692 | 4158 | 3890 |
| 2 | medRT | ML | 1 | Intercept (VC) | None | 135 | 4289 | 3751 | 3481 |
| 3 | medRT | ML | 1 | Intercept + Visit (VC) | None | 136 | 4075 | 3533 | 3261 |
| 4 | medRT | ML | 1 | Intercept + Type (VC) | None | 136 | 4294 | 3753 | 3481 |
| 5 | medRT | ML | 1 | Intercept + Visit + Type (VC) | None | 137 | 4027 | 3481 | 3207 |
| 6 | medRT | ML | 1 | Intercept + Visit + Type (VC) | Visit (ID*Type) (AR1) | 138 | 4033 | 3483 | 3207 |
| *7 | medRT | ML | 1 | Intercept + Visit + Type (VC) | Visit (ID*Type) (CS) | 138 | 4224 | 3675 | 3399 |
| *8 | medRT | ML | 1 | Intercept + Visit + Type (VC) | Visit (ID*Type) (SP_POWER) (t) | 138 | 4033 | 3483 | 3207 |
| ^9 | medRT | ML | 1 | Intercept + Visit + Type (VC) | Visit (ID*Type) (ARMA11) | 139 | 4038 | 3485 | 3207 |
| 10 | medRT | ML | 1 | Intercept + Visit + Type (VC) | Visit (ID*Type) (ARH1) | 148 | 4067 | 3478 | 3182 |
| *11 | medRT | ML | 1 | Intercept + Visit + Type (VC) | Visit (ID*Type) (CSH) | 148 | 4105 | 3516 | 3220 |
| 12 | medRT | REML | 1 | Intercept + Visit + Type (VC) | Visit (ID*Type) (ARH1) | 148 | 2453 | 2400 | 2370 |
| Final | medRT | REML | 1 + $Iz_{between}$ $Iz_{between}$*Group $Iz_{between}$*Sex $Iz_{within}$ $Iz_{within}$*Group $Iz_{within}$*Sex | Intercept + Visit + Type (VC) | Visit (ID*Type) (ARH1) | 154 | 2399 | 2346 | 2316 |

*Failed to converge; ^Hessian matrix not positive definite

| Model | Data | Method | Fixed (FE) | Random (Cov) | Repeated (Cov) | Param | BIC | AIC | -2 LL |
|---|---|---|---|---|---|---|---|---|---|
| 1 | Errors | ML | Age Group Sex Visit Type Group*Sex Group*Visit Group*Sex*Visit Group*Type Group*Sex*Type Group*Sex*Type*Visit | None | None | 134 | 1954 | 1420 | 1152 |
| 2 | Errors | ML | 1 | Intercept (VC) | None | 135 | 1891 | 1354 | 1084 |
| *3 | Errors | ML | 1 | Intercept + Visit (VC) | None | 136 | 1897 | 1356 | 1084 |
| 4 | Errors | ML | 1 | Intercept + Type (VC) | None | 136 | 1819 | 1278 | 1006 |
| *5 | Errors | ML | 1 | Intercept + Visit + Type (VC) | None | 137 | 1825 | 1280 | 1006 |
| 6 | Errors | ML | 1 | Intercept + Type (VC) | Visit (ID*Type) (AR1) | 137 | 1823 | 1277 | 1003 |
| 7 | Errors | ML | 1 | Intercept + Type (VC) | Visit (ID*Type) (CS) | 137 | 1825 | 1280 | 1006 |
| *8 | Errors | ML | 1 | Intercept + Type (VC) | Visit (ID*Type) (SP_POWER) (t) | 137 | 1888 | 1342 | 1068 |
| *9 | Errors | ML | 1 | Intercept + Type (VC) | Visit (ID*Type) (ARMA11) | 138 | 1829 | 1280 | 1004 |
| 10 | Errors | ML | 1 | Intercept + Type (VC) | Visit (ID*Type) (ARH1) | 147 | 1865 | 1280 | 986 |
| *11 | Errors | ML | 1 | Intercept + Type (VC) | Visit (ID*Type) (CSH) | 147 | 1401 | 816 | 522 |
| 12 | Errors | REML | 1 | Intercept + Type (VC) | None | 136 | 943 | 933 | 927 |
| *Final | Errors | REML | 1 + $Iz_{between}$ $Iz_{between}$*Group $Iz_{between}$*Sex $Iz_{within}$ $Iz_{within}$*Group $Iz_{within}$*Sex | Intercept + Type (VC) | None | 142 | 937 | 926 | 920 |

*Failed to converge

| Model | Data | Method | Fixed (FE) | Random (Cov) | Repeated (Cov) | Param | BIC | AIC | -2 LL |
|---|---|---|---|---|---|---|---|---|---|
| 1 | Lapses | ML | Age Group Sex Visit Type Group*Sex Group*Visit Group*Sex*Visit Group*Type Group*Sex*Type Group*Sex*Type*Visit | None | None | 134 | 1683 | 1150 | 882 |
| 2 | Lapses | ML | 1 | Intercept (VC) | None | 135 | 1683 | 1145 | 875 |
| 3 | Lapses | ML | 1 | Intercept + Visit (VC) | None | 136 | 1674 | 1132 | 860 |
| 4 | Lapses | ML | 1 | Intercept + Type (VC) | None | 136 | 1689 | 1147 | 875 |
| 5 | Lapses | ML | 1 | Intercept + Visit + Type (VC) | None | 137 | 1679 | 1133 | 859 |
| 6 | Lapses | ML | 1 | Intercept + Visit (VC) | Visit (ID*Type) (AR1) | 137 | 1679 | 1134 | 860 |
| 7 | Lapses | ML | 1 | Intercept + Visit (VC) | Visit (ID*Type) (CS) | 137 | 1679 | 1133 | 859 |
| 8 | Lapses | ML | 1 | Intercept + Visit (VC) | Visit (ID*Type) (SP_POWER) (t) | 137 | 1679 | 1134 | 860 |
| 9 | Lapses | ML | 1 | Intercept + Visit (VC) | Visit (ID*Type) (ARMA11) | 138 | 1685 | 1136 | 860 |
| 10 | Lapses | ML | 1 | Intercept + Visit (VC) | Visit (ID*Type) (ARH1) | 147 | 1688 | 1103 | 809 |
| 11 | Lapses | ML | 1 | Intercept + Visit (VC) | Visit (ID*Type) (CSH) | 147 | 1688 | 1102 | 808 |
| 12 | Lapses | REML | 1 | Intercept + Visit (VC) | Visit (ID*Type) (ARH1) | 147 | 875 | 825 | 797 |
| Final | Lapses | REML | 1 + $Iz_{between}$ $Iz_{between}$*Group $Iz_{between}$*Sex $Iz_{within}$ $Iz_{within}$*Group $Iz_{within}$*Sex | Intercept + Visit (VC) | Visit (ID*Type) (ARH1) | 153 | 878 | 828 | 800 |

| Model | Data | Method | Fixed (FE) | Random (Cov) | Repeated (Cov) | Param | BIC | AIC | -2 LL |
|---|---|---|---|---|---|---|---|---|---|
| 1 | Rejects | ML | Age Group Sex Visit Type Group*Sex Group*Visit Group*Sex*Visit Group*Type Group*Sex*Type Group*Sex*Type*Visit | None | None | 134 | 2124 | 1591 | 1323 |
| 2 | Rejects | ML | 1 | Intercept (VC) | None | 135 | 2066 | 1528 | 1258 |
| *3 | Rejects | ML | 1 | Intercept + Visit (VC) | None | 136 | 2072 | 1530 | 1258 |
| 4 | Rejects | ML | 1 | Intercept + Type (VC) | None | 136 | 1999 | 1458 | 1186 |
| 5 | Rejects | ML | 1 | Intercept + Visit + Type (VC) | None | 137 | 2000 | 1455 | 1181 |
| 6 | Rejects | ML | 1 | Intercept + Visit + Type (VC) | Visit (ID*Type) (AR1) | 138 | 2006 | 1457 | 1181 |
| ^7 | Rejects | ML | 1 | Intercept + Visit + Type (VC) | Visit (ID*Type) (CS) | 138 | 2006 | 1457 | 1181 |
| *8 | Rejects | ML | 1 | Intercept + Visit + Type (VC) | Visit (ID*Type) (SP_POWER) (t) | 138 | 2094 | 1545 | 1269 |
| *9 | Rejects | ML | 1 | Intercept + Visit + Type (VC) | Visit (ID*Type) (ARMA11) | 139 | 1559 | 1006 | 728 |
| 10 | Rejects | ML | 1 | Intercept + Visit + Type (VC) | Visit (ID*Type) (ARH1) | 148 | 2033 | 1444 | 1148 |
| *11 | Rejects | ML | 1 | Intercept + Visit + Type (VC) | Visit (ID*Type) (CSH) | 148 | 2051 | 1462 | 1166 |
| 12 | Rejects | REML | 1 | Intercept + Visit + Type (VC) | Visit (ID*Type) (ARH1) | 148 | 1105 | 1051 | 1021 |
| ^Final | Rejects | REML | 1 + $Iz_{between}$ $Iz_{between}$*Group $Iz_{between}$*Sex $Iz_{within}$ $Iz_{within}$*Group $Iz_{within}$*Sex | Intercept + Visit + Type (VC) | Visit (ID*Type) (ARH1) | 154 | 1096 | 1043 | 1013 |

*Failed to converge; ^Hessian matrix not positive definite

#### B. Wisconsin Card Sorting Test

| Model | Data | Method | Fixed (FE) | Random (Cov) | Repeated (Cov) | Param | BIC | AIC | -2 LL |
|---|---|---|---|---|---|---|---|---|---|
| 1 | medRT | ML | Age Group Sex Visit Group*Sex Group*Visit Group*Sex*Visit | None | None | 46 | 1774 | 1642 | 1550 |
| 2 | medRT | ML | 1 | Intercept (VC) | None | 47 | 1657 | 1522 | 1428 |
| ^3 | medRT | ML | 1 | Intercept + Visit (VC) | None | 48 | 1662 | 1524 | 1428 |
| 4 | medRT | ML | 1 | Intercept (VC) | Visit (ID) (AR1) | 48 | 1630 | 1491 | 1395 |
| *5 | medRT | ML | 1 | Intercept (VC) | Visit (ID) (CS) | 48 | 1662 | 1524 | 1428 |
| 6 | medRT | ML | 1 | Intercept (VC) | Visit (ID) (SP_POWER) (t) | 48 | 1636 | 1497 | 1401 |
| 7 | medRT | ML | 1 | Intercept (VC) | Visit (ID) (ARMA11) | 49 | 1632 | 1491 | 1393 |
| ^8 | medRT | ML | 1 | Intercept (VC) | Visit (ID) (ARH1) | 58 | -65968 | -66135 | -66251 |
| 9 | medRT | ML | 1 | Intercept (VC) | Visit (ID) (CSH) | 58 | 1671 | 1504 | 1388 |
| 10 | medRT | REML | 1 | Intercept (VC) | Visit (ID) (AR1) | 48 | 1022 | 1015 | 1009 |
| Final | medRT | REML | 1 + $Iz_{between}$ $Iz_{between}$*Group $Iz_{between}$*Sex $Iz_{within}$ $Iz_{within}$*Group $Iz_{within}$*Sex | Intercept (VC) | Visit (ID) (AR1) | 54 | 963 | 956 | 950 |

*Failed to converge; ^Hessian matrix not positive definite

| Model | Data | Method | Fixed (FE) | Random (Cov) | Repeated (Cov) | Param | BIC | AIC | -2 LL |
|---|---|---|---|---|---|---|---|---|---|
| 1 | Errors | ML | Age Group Sex Visit Type Group*Sex Group*Visit Group*Sex*Visit Group*Type Group*Sex*Type Group*Sex*Type*Visit | None | None | 90 | 1727 | 1405 | 1225 |
| 2 | Errors | ML | 1 | Intercept (VC) | None | 91 | 1719 | 1394 | 1212 |
| 3 | Errors | ML | 1 | Intercept + Visit (VC) | None | 92 | 1725 | 1396 | 1212 |
| 4 | Errors | ML | 1 | Intercept + Type (VC) | None | 92 | 1717 | 1388 | 1204 |
| 5 | Errors | ML | 1 | Intercept + Visit + Type (VC) | None | 93 | 1721 | 1388 | 1202 |
| 6 | Errors | ML | 1 | Intercept + Type (VC) | Visit (ID*Type) (AR1) | 93 | 1720 | 1387 | 1201 |
| ^7 | Errors | ML | 1 | Intercept + Type (VC) | Visit (ID*Type) (CS) | 93 | 1723 | 1390 | 1204 |
| 8 | Errors | ML | 1 | Intercept + Type (VC) | Visit (ID*Type) (SP_POWER) (t) | 93 | 1719 | 1386 | 1200 |
| ^9 | Errors | ML | 1 | Intercept + Type (VC) | Visit (ID*Type) (ARMA11) | 94 | 1726 | 1390 | 1202 |
| 10 | Errors | ML | 1 | Intercept + Type (VC) | Visit (ID*Type) (ARH1) | 103 | 1748 | 1379 | 1173 |
| *11 | Errors | ML | 1 | Intercept + Type (VC) | Visit (ID*Type) (CSH) | 103 | 1808 | 1439 | 1233 |
| 12 | Errors | REML | 1 | Intercept + Type (VC) | None | 92 | 989 | 980 | 974 |
| Final | Errors | REML | 1 + $Iz_{between}$ $Iz_{between}$*Group $Iz_{between}$*Sex $Iz_{within}$ $Iz_{within}$*Group $Iz_{within}$*Sex | Intercept + Type (VC) | None | 98 | 982 | 973 | 967 |

^Hessian matrix not positive definite; *Failed to converge

| Model | Data | Method | Fixed (FE) | Random (Cov) | Repeated (Cov) | Param | BIC | AIC | -2 LL |
|---|---|---|---|---|---|---|---|---|---|
| 1 | Lapses | ML | Age Group Sex Visit Type Group*Sex Group*Visit Group*Sex*Visit Group*Type Group*Sex*Type Group*Sex*Type*Visit | None | None | 90 | 1622 | 1300 | 1120 |
| 2 | Lapses | ML | 1 | Intercept (VC) | None | 91 | 1625 | 1299 | 1117 |
| 3 | Lapses | ML | 1 | Intercept + Visit (VC) | None | 92 | 1624 | 1295 | 1111 |
| 4 | Lapses | ML | 1 | Intercept + Type (VC) | None | 92 | 1629 | 1300 | 1116 |
| 5 | Lapses | ML | 1 | Intercept + Visit + Type (VC) | None | 93 | 1626 | 1294 | 1108 |
| 6 | Lapses | ML | 1 | Intercept + Visit (VC) | Visit (ID*Type) (AR1) | 93 | 1628 | 1296 | 1110 |
| 7 | Lapses | ML | 1 | Intercept + Visit (VC) | Visit (ID*Type) (CS) | 93 | 1626 | 1294 | 1108 |
| 8 | Lapses | ML | 1 | Intercept + Visit (VC) | Visit (ID*Type) (SP_POWER) (t) | 93 | 1628 | 1296 | 1110 |
| 9 | Lapses | ML | 1 | Intercept + Visit (VC) | Visit (ID*Type) (ARMA11) | 94 | 1630 | 1294 | 1106 |
| 10 | Lapses | ML | 1 | Intercept + Visit (VC) | Visit (ID*Type) (ARH1) | 103 | 1595 | 1226 | 1020 |
| 11 | Lapses | ML | 1 | Intercept + Visit (VC) | Visit (ID*Type) (CSH) | 103 | 1595 | 1227 | 1021 |
| 12 | Lapses | REML | 1 | Intercept + Visit (VC) | Visit (ID*Type) (ARH1) | 103 | 925 | 881 | 853 |
| Final | Lapses | REML | 1 + $Iz_{between}$ $Iz_{between}$*Group $Iz_{between}$*Sex $Iz_{within}$ $Iz_{within}$*Group $Iz_{within}$*Sex | Intercept + Visit (VC) | Visit (ID*Type) (ARH1) | 109 | 917 | 873 | 845 |

| Model | Data | Method | Fixed (FE) | Random (Cov) | Repeated (Cov) | Param | BIC | AIC | -2 LL |
|---|---|---|---|---|---|---|---|---|---|
| 1 | Rejects | ML | Age Group Sex Visit Type Group*Sex Group*Visit Group*Sex*Visit Group*Type Group*Sex*Type Group*Sex*Type*Visit | None | None | 90 | 1766 | 1445 | 1265 |
| 2 | Rejects | ML | 1 | Intercept (VC) | None | 91 | 1765 | 1440 | 1258 |
| *3 | Rejects | ML | 1 | Intercept + Visit (VC) | None | 92 | 1771 | 1442 | 1258 |
| *4 | Rejects | ML | 1 | Intercept + Type (VC) | None | 92 | 1741 | 1412 | 1228 |
| *5 | Rejects | ML | 1 | Intercept + Visit + Type (VC) | None | 93 | 1745 | 1412 | 1226 |
| 6 | Rejects | ML | 1 | Intercept (VC) | Visit (ID*Type) (AR1) | 92 | 1768 | 1439 | 1255 |
| ^7 | Rejects | ML | 1 | Intercept (VC) | Visit (ID*Type) (CS) | 92 | 1741 | 1412 | 1228 |
| 8 | Rejects | ML | 1 | Intercept (VC) | Visit (ID*Type) (SP_POWER) (t) | 92 | 1769 | 1440 | 1256 |
| ^9 | Rejects | ML | 1 | Intercept (VC) | Visit (ID*Type) (ARMA11) | 93 | 1746 | 1414 | 1228 |
| 10 | Rejects | ML | 1 | Intercept (VC) | Visit (ID*Type) (ARH1) | 102 | 1811 | 1446 | 1242 |
| ^11 | Rejects | ML | 1 | Intercept (VC) | Visit (ID*Type) (CSH) | 102 | 1785 | 1420 | 1216 |
| 12 | Rejects | REML | 1 | Intercept (VC) | None | 91 | 1020 | 1014 | 1010 |
| Final | Rejects | REML | 1 + $Iz_{between}$ $Iz_{between}$*Group $Iz_{between}$*Sex $Iz_{within}$ $Iz_{within}$*Group $Iz_{within}$*Sex | Intercept (VC) | None | 97 | 1009 | 1002 | 998 |

^Hessian matrix not positive definite; *Failed to converge

#### C. Sternberg Working Memory Task

| Model | Data | Method | Fixed (FE) | Random (Cov) | Repeated (Cov) | Param | BIC | AIC | -2 LL |
|---|---|---|---|---|---|---|---|---|---|
| 1 | medRT | ML | Age Group Sex Visit Type Group*Sex Group*Visit Group*Sex*Visit Group*Type Group*Sex*Type Group*Sex*Type*Visit | None | None | 134 | 5579 | 5045 | 4777 |
| 2 | medRT | ML | 1 | Intercept (VC) | None | 135 | 5451 | 4913 | 4643 |
| 3 | medRT | ML | 1 | Intercept + Visit (VC) | None | 136 | 5456 | 4915 | 4643 |
| 4 | medRT | ML | 1 | Intercept + Type (VC) | None | 136 | 5404 | 4863 | 4591 |
| 5 | medRT | ML | 1 | Intercept + Visit + Type (VC) | None | 137 | 5404 | 4858 | 4584 |
| 6 | medRT | ML | 1 | Intercept + Visit + Type (VC) | Visit (ID*Type) (AR1) | 138 | 5402 | 4853 | 4577 |
| *7 | medRT | ML | 1 | Intercept + Visit + Type (VC) | Visit (ID*Type) (CS) | 138 | 5410 | 4860 | 4584 |
| 8 | medRT | ML | 1 | Intercept + Visit + Type (VC) | Visit (ID*Type) (SP_POWER) (t) | 138 | 5404 | 4855 | 4579 |
| 9 | medRT | ML | 1 | Intercept + Visit + Type (VC) | Visit (ID*Type) (ARMA11) | 139 | 5408 | 4854 | 4576 |
| 10 | medRT | ML | 1 | Intercept + Visit + Type (VC) | Visit (ID*Type) (ARH1) | 148 | 5441 | 4852 | 4556 |
| *11 | medRT | ML | 1 | Intercept + Visit + Type (VC) | Visit (ID*Type) (CSH) | 148 | 5471 | 4881 | 4585 |
| 12 | medRT | REML | 1 | Intercept + Visit + Type (VC) | Visit (ID*Type) (AR1) | 138 | 3326 | 3308 | 3298 |
| Final | medRT | REML | 1 + $Iz_{between}$ $Iz_{between}$*Group $Iz_{between}$*Sex $Iz_{within}$ $Iz_{within}$*Group $Iz_{within}$*Sex | Intercept + Visit + Type (VC) | Visit (ID*Type) (AR1) | 144 | 3269 | 3251 | 3241 |

*Failed to converge

| Model | Data | Method | Fixed (FE) | Random (Cov) | Repeated (Cov) | Param | BIC | AIC | -2 LL |
|---|---|---|---|---|---|---|---|---|---|
| 1 | Errors | ML | Age Group Sex Visit Type Group*Sex Group*Visit Group*Sex*Visit Group*Type Group*Sex*Type Group*Sex*Type*Visit | None | None | 134 | 1810 | 1276 | 1008 |
| 2 | Errors | ML | 1 | Intercept (VC) | None | 135 | 1756 | 1219 | 949 |
| *3 | Errors | ML | 1 | Intercept + Visit (VC) | None | 136 | 1762 | 1221 | 949 |
| 4 | Errors | ML | 1 | Intercept + Type (VC) | None | 136 | 1740 | 1198 | 926 |
| *5 | Errors | ML | 1 | Intercept + Visit + Type (VC) | None | 137 | 1745 | 1200 | 926 |
| 6 | Errors | ML | 1 | Intercept + Type (VC) | Visit (ID*Type) (AR1) | 137 | 1744 | 1198 | 924 |
| *7 | Errors | ML | 1 | Intercept + Type (VC) | Visit (ID*Type) (CS) | 137 | 1749 | 1203 | 929 |
| 8 | Errors | ML | 1 | Intercept + Type (VC) | Visit (ID*Type) (SP_POWER) (t) | 137 | 1742 | 1196 | 922 |
| 9 | Errors | ML | 1 | Intercept + Type (VC) | Visit (ID*Type) (ARMA11) | 138 | 1747 | 1197 | 921 |
| 10 | Errors | ML | 1 | Intercept + Type (VC) | Visit (ID*Type) (ARH1) | 147 | 1786 | 1201 | 907 |
| *11 | Errors | ML | 1 | Intercept + Type (VC) | Visit (ID*Type) (CSH) | 147 | 1607 | 1022 | 728 |
| 12 | Errors | REML | 1 | Intercept + Type (VC) | Visit (ID*Type) (SP_POWER) (t) | 137 | 894 | 880 | 872 |
| Final | Errors | REML | 1 + $Iz_{between}$ $Iz_{between}$*Group $Iz_{between}$*Sex $Iz_{within}$ $Iz_{within}$*Group $Iz_{within}$*Sex | Intercept + Type (VC) | Visit (ID*Type) (SP_POWER) (t) | 143 | 897 | 883 | 875 |

*Failed to converge

| Model | Data | Method | Fixed (FE) | Random (Cov) | Repeated (Cov) | Param | BIC | AIC | -2 LL |
|---|---|---|---|---|---|---|---|---|---|
| 1 | Lapses | ML | Age Group Sex Visit Type Group*Sex Group*Visit Group*Sex*Visit Group*Type Group*Sex*Type Group*Sex*Type*Visit | None | None | 134 | 1947 | 1414 | 1146 |
| 2 | Lapses | ML | 1 | Intercept (VC) | None | 135 | 1907 | 1369 | 1099 |
| *3 | Lapses | ML | 1 | Intercept + Visit (VC) | None | 136 | 1913 | 1371 | 1099 |
| 4 | Lapses | ML | 1 | Intercept + Type (VC) | None | 136 | 1872 | 1330 | 1058 |
| *5 | Lapses | ML | 1 | Intercept + Visit + Type (VC) | None | 137 | 1878 | 1332 | 1058 |
| 6 | Lapses | ML | 1 | Intercept + Type (VC) | Visit (ID*Type) (AR1) | 137 | 1875 | 1330 | 1056 |
| 7 | Lapses | ML | 1 | Intercept + Type (VC) | Visit (ID*Type) (CS) | 137 | 1878 | 1332 | 1058 |
| *8 | Lapses | ML | 1 | Intercept + Type (VC) | Visit (ID*Type) (SP_POWER) (t) | 137 | 2002 | 1457 | 1183 |
| 9 | Lapses | ML | 1 | Intercept + Type (VC) | Visit (ID*Type) (ARMA11) | 138 | 1881 | 1332 | 1056 |
| 10 | Lapses | ML | 1 | Intercept + Type (VC) | Visit (ID*Type) (ARH1) | 147 | 1882 | 1297 | 1003 |
| *11 | Lapses | ML | 1 | Intercept + Type (VC) | Visit (ID*Type) (CSH) | 147 | 1607 | 1022 | 728 |
| 12 | Lapses | REML | 1 | Intercept + Type (VC) | None | 136 | 979 | 968 | 962 |
| Final | Lapses | REML | 1 + $Iz_{between}$ $Iz_{between}$*Group $Iz_{between}$*Sex $Iz_{within}$ $Iz_{within}$*Group $Iz_{within}$*Sex | Intercept + Type (VC) | None | 142 | 977 | 967 | 961 |

*Failed to converge

| Model | Data | Method | Fixed (FE) | Random (Cov) | Repeated (Cov) | Param | BIC | AIC | -2 LL |
|---|---|---|---|---|---|---|---|---|---|
| 1 | Rejects | ML | Age Group Sex Visit Type Group*Sex Group*Visit Group*Sex*Visit Group*Type Group*Sex*Type Group*Sex*Type*Visit | None | None | 134 | 2074 | 1540 | 1272 |
| 2 | Rejects | ML | 1 | Intercept (VC) | None | 135 | 2070 | 1532 | 1262 |
| *3 | Rejects | ML | 1 | Intercept + Visit (VC) | None | 136 | 2075 | 1534 | 1262 |
| 4 | Rejects | ML | 1 | Intercept + Type (VC) | None | 136 | 2066 | 1525 | 1253 |
| *5 | Rejects | ML | 1 | Intercept + Visit + Type (VC) | None | 137 | 2072 | 1527 | 1253 |
| 6 | Rejects | ML | 1 | Intercept + Type (VC) | Visit (ID*Type) (AR1) | 137 | 2071 | 1525 | 1251 |
| ^7 | Rejects | ML | 1 | Intercept + Type (VC) | Visit (ID*Type) (CS) | 137 | 2072 | 1527 | 1253 |
| *8 | Rejects | ML | 1 | Intercept + Type (VC) | Visit (ID*Type) (SP_POWER) (t) | 137 | 2324 | 1779 | 1505 |
| 9 | Rejects | ML | 1 | Intercept + Type (VC) | Visit (ID*Type) (ARMA11) | 138 | 2075 | 1526 | 1250 |
| 10 | Rejects | ML | 1 | Intercept + Type (VC) | Visit (ID*Type) (ARH1) | 147 | 2077 | 1492 | 1198 |
| *11 | Rejects | ML | 1 | Intercept + Type (VC) | Visit (ID*Type) (CSH) | 147 | 1607 | 1022 | 728 |
| 12 | Rejects | REML | 1 | Intercept + Type (VC) | Visit (ID*Type) (ARH1) | 147 | 1134 | 1084 | 1056 |
| ^Final | Rejects | REML | 1 + $Iz_{between}$ $Iz_{between}$*Group $Iz_{between}$*Sex $Iz_{within}$ $Iz_{within}$*Group $Iz_{within}$*Sex | Intercept + Type (VC) | Visit (ID*Type) (ARH1) | 153 | 1129 | 1080 | 1052 |

*Failed to converge; ^Hessian matrix not positive definite

| Model | Data | Method | Fixed (FE) | Random (Cov) | Repeated (Cov) | Param | BIC | AIC | -2 LL |
|---|---|---|---|---|---|---|---|---|---|
| 1 | TEPR | ML | Age Group Sex Visit Task Type Group*Sex Group*Visit Group*Task Group*Type Sex*Visit Sex*Task Sex*Type Visit*Task Visit*Type Task*Type Group*Sex*Visit Group*Sex*Task Group*Sex*Type Group*Visit*Task Group*Visit*Type Sex*Visit*Task Sex*Visit*Type Group*Sex*Visit*Task Group*Sex*Visit*Type Group*Sex*Task*Type Group*Visit*Task*Type Sex*Visit*Task*Type | None | None | 246 | 1813 | 633 | 171 |
| 2 | TEPR | ML | 1 | Intercept (VC) | None | 247 | 1026 | -128 | -622 |
| 3 | TEPR | ML | 1 | Intercept + Visit (VC) | None | 248 | 290 | -869 | -1365 |
| 4 | TEPR | ML | 1 | Intercept + Task (VC) | None | 248 | 1017 | -142 | -638 |
| 5 | TEPR | ML | 1 | Intercept + Type (VC) | None | 248 | 1033 | -126 | -622 |
| 6 | TEPR | ML | 1 | Intercept + Visit + Task (VC) | None | 249 | 143 | -1021 | -1519 |
| 7 | TEPR | ML | 1 | Intercept + Visit + Type (VC) | None | 249 | 292 | -872 | -1370 |
| 8 | TEPR | ML | 1 | Intercept + Visit + Task + Type (VC) | None | 250 | 137 | -1031 | -1531 |
| 9 | TEPR | ML | 1 | Intercept + Visit + Task +Type (VC) | Visit (ID*TaskType) (AR1) | 251 | 144 | -1029 | -1531 |
| 10 | TEPR | ML | 1 | Intercept + Visit + Task +Type (VC) | Visit (ID*TaskType) (CS) | 251 | 144 | -1029 | -1531 |
| 11 | TEPR | ML | 1 | Intercept + Visit + Task +Type (VC) | Visit (ID*TaskType) (CSH) | 261 | 199 | -1021 | -1543 |
| 12 | TEPR | ML | 1 | Intercept + Visit + Task +Type (VC) | Visit (ID*TaskType) (ARH1) | 261 | 199 | -1021 | -1543 |
| 13 | TEPR | ML | 1 | Intercept + Visit + Task +Type (VC) | Visit (ID*TaskType) (SP_POWER) (t) | 251 | 144 | -1030 | -1532 |
| 14 | TEPR | REML | 1 | Intercept + Visit + Task + Type (VC) | None | 250 | -509 | -530 | -540 |
| Final | TEPR | REML | 1 + $Iz_{between}$ $Iz_{between}$*Group $Iz_{between}$*Sex $Iz_{within}$ $Iz_{within}$*Group $Iz_{within}$*Sex | Intercept + Visit + Task + Type (VC) | None | 256 | -496 | -517 | -527 |

#### D. Resting Tonic Physiological measures

| Model | Data | Method | Fixed (FE) | Random (Cov) | Repeated (Cov) | Param | BIC | AIC | -2 LL |
|---|---|---|---|---|---|---|---|---|---|
| 1 | mPupil | ML | Age Group Sex Visit TP Visit*TP Group*Sex Group*Visit Group*Sex*Visit Group*TP Group*Sex*TP Group*Sex*Visit*TP | None | None | 134 | 1359 | 825 | 557 |
| 2 | mPupil | ML | 1 | Intercept (VC) | None | 135 | 854 | 316 | 46 |
| 3 | mPupil | ML | 1 | Intercept + Visit (VC) | None | 136 | 666 | 124 | -148 |
| 4 | mPupil | ML | 1 | Intercept + TP (VC) | None | 136 | 859 | 317 | 45 |
| 5 | mPupil | ML | 1 | Intercept + Visit + TP (VC) | None | 137 | 617 | 72 | -202 |
| 6 | mPupil | ML | 1 | Intercept + Visit + TP (VC) | Visit (ID*TP) (AR1) | 138 | 622 | 72 | -204 |
| *7 | mPupil | ML | 1 | Intercept + Visit + TP (VC) | Visit (ID*TP) (CS) | 138 | 1553 | 1003 | 728 |
| 8 | mPupil | ML | 1 | Intercept + Visit + TP (VC) | Visit (ID*TP) (SP_POWER) (t) | 138 | 623 | 73 | -203 |
| 9 | mPupil | ML | 1 | Intercept + Visit + TP (VC) | Visit (ID*TP) (ARMA11) | 139 | 626 | 72 | -206 |
| 10 | mPupil | ML | 1 | Intercept + Visit + TP (VC) | Visit (ID*TP) (DIAG) | 147 | 640 | 54 | -240 |
| 11 | mPupil | ML | 1 | Intercept + Visit + TP (VC) | Visit (ID*TP) (ARH1) | 148 | 645 | 55 | -241 |
| *12 | mPupil | ML | 1 | Intercept + Visit + TP (VC) | Visit (ID*TP) (CSH) | 148 | 110 | -479 | -775 |
| 13 | mPupil | REML | 1 | Intercept + Visit + TP (VC) | Visit (ID*TP) (DIAG) | 147 | 175 | 125 | 97 |
| Final | mPupil | REML | 1 + $Iz_{between}$ $Iz_{between}$*Group $Iz_{between}$*Sex $Iz_{within}$ $Iz_{within}$*Group $Iz_{within}$*Sex | Intercept + Visit + TP (VC) | Visit (ID*TP) (DIAG) | 153 | 181 | 132 | 104 |

*Failed to converge; TP = resting assessment time point (Pre, Stim, Post)

| Model | Data | Method | Fixed (FE) | Random (Cov) | Repeated (Cov) | Param | BIC | AIC | -2 LL |
|---|---|---|---|---|---|---|---|---|---|
| 1 | mSCL | ML | Age Group Sex Visit TP Visit*TP Group*Sex Group*Visit Group*Sex*Visit Group*TP Group*Sex*TP Group*Sex*Visit*TP | None | None | 134 | 3051 | 2518 | 2250 |
| 2 | mSCL | ML | 1 | Intercept (VC) | None | 135 | 2911 | 2374 | 2104 |
| 3 | mSCL | ML | 1 | Intercept + Visit (VC) | None | 136 | 2737 | 2195 | 1923 |
| 4 | mSCL | ML | 1 | Intercept + TP (VC) | None | 136 | 2915 | 2373 | 2101 |
| 5 | mSCL | ML | 1 | Intercept + Visit + TP (VC) | None | 137 | 2676 | 2130 | 1856 |
| 6 | mSCL | ML | 1 | Intercept + Visit + TP (VC) | Visit (ID*TP) (AR1) | 138 | 2680 | 2130 | 1854 |
| 7 | mSCL | ML | 1 | Intercept + Visit + TP (VC) | Visit (ID*TP) (CS) | 138 | 2682 | 2132 | 1856 |
| 8 | mSCL | ML | 1 | Intercept + Visit + TP (VC) | Visit (ID*TP) (SP_POWER) (t) | 138 | 2679 | 2129 | 1853 |
| *9 | mSCL | ML | 1 | Intercept + Visit + TP (VC) | Visit (ID*TP) (ARMA11) | 139 | 2686 | 2132 | 1854 |
| 10 | mSCL | ML | 1 | Intercept + Visit + TP (VC) | Visit (ID*TP) (DIAG) | 147 | 2720 | 2135 | 1841 |
| 11 | mSCL | ML | 1 | Intercept + Visit + TP (VC) | Visit (ID*TP) (ARH1) | 148 | 2723 | 2134 | 1838 |
| *12 | mSCL | ML | 1 | Intercept + Visit + TP (VC) | Visit (ID*TP) (CSH) | 148 | 2723 | 2134 | 1838 |
| 13 | mSCL | REML | 1 | Intercept + Visit + TP (VC) | None | 137 | 1513 | 1499 | 1491 |
| Final | mSCL | REML | 1 + $Iz_{between}$ $Iz_{between}$*Group $Iz_{between}$*Sex $Iz_{within}$ $Iz_{within}$*Group $Iz_{within}$*Sex | Intercept + Visit + TP (VC) | None | 143 | 1488 | 1474 | 1466 |

*Failed to converge; ^Hessian matrix not positive definite; TP = resting assessment time point (Pre, Stim, Post)

| Model | Data | Method | Fixed (FE) | Random (Cov) | Repeated (Cov) | Param | BIC | AIC | -2 LL |
|---|---|---|---|---|---|---|---|---|---|
| 1 | mRR | ML | Age Group Sex Visit TP Visit*TP Group*Sex Group*Visit Group*Sex*Visit Group*TP Group*Sex*TP Group*Sex*Visit*TP | None | None | 134 | 5547 | 5014 | 4746 |
| 2 | mRR | ML | 1 | Intercept (VC) | None | 135 | 5059 | 4522 | 4252 |
| 3 | mRR | ML | 1 | Intercept + Visit (VC) | None | 136 | 4682 | 4141 | 3869 |
| *4 | mRR | ML | 1 | Intercept + TP (VC) | None | 136 | 5065 | 4524 | 4252 |
| 5 | mRR | ML | 1 | Intercept + Visit + TP (VC) | None | 137 | 4646 | 4100 | 3826 |
| 6 | mRR | ML | 1 | Intercept + Visit + TP (VC) | Visit (ID*TP) (AR1) | 138 | 4651 | 4102 | 3826 |
| *7 | mRR | ML | 1 | Intercept + Visit + TP (VC) | Visit (ID*TP) (CS) | 138 | 5013 | 4464 | 4188 |
| *8 | mRR | ML | 1 | Intercept + Visit + TP (VC) | Visit (ID*TP) (SP_POWER) (t) | 138 | 4652 | 4102 | 3826 |
| ^10 | mRR | ML | 1 | Intercept + Visit + TP (VC) | Visit (ID*TP) (ARMA11) | 139 | 4658 | 4104 | 3826 |

| | | | | | | | | | |
|---|---|---|---|---|---|---|---|---|---|
| 9 | mRR | ML | 1 | Intercept + Visit + TP (VC) | Visit (ID*TP) (DIAG) | 147 | 4654 | 4069 | 3775 |
| 11 | mRR | ML | 1 | Intercept + Visit + TP (VC) | Visit (ID*TP) (ARH1) | 148 | 4660 | 4071 | 3775 |
| *12 | mRR | ML | 1 | Intercept + Visit + TP (VC) | Visit (ID*TP) (CSH) | 148 | 4700 | 4111 | 3815 |
| 13 | mRR | REML | 1 | Intercept + Visit + TP (VC) | Visit (ID*TP) (DIAG) | 147 | 2841 | 2791 | 2763 |
| Final | mRR | REML | 1 + $Iz_{between}$ $Iz_{between}$*Group $Iz_{between}$*Sex $Iz_{within}$ $Iz_{within}$*Group $Iz_{within}$*Sex | Intercept + Visit + TP (VC) | Visit (ID*TP) (DIAG) | 153 | 2781 | 2731 | 2704 |

*Failed to converge; ^Hessian matrix not positive definite; TP = resting assessment time point (Pre, Stim, Post)

| Model | Data | Method | Fixed (FE) | Random (Cov) | Repeated (Cov) | Param | BIC | AIC | -2 LL |
|---|---|---|---|---|---|---|---|---|---|
| 1 | mResp | ML | Age Group Sex Visit TP Visit*TP Group*Sex Group*Visit Group*Sex*Visit Group*TP Group*Sex*TP Group*Sex*Visit*TP | None | None | 134 | -623 | -1156 | -1424 |
| 2 | mResp | ML | 1 | Intercept (VC) | None | 135 | -837 | -1375 | -1645 |
| 3 | mResp | ML | 1 | Intercept + Visit (VC) | None | 136 | -998 | -1540 | -1812 |
| 4 | mResp | ML | 1 | Intercept + TP (VC) | None | 136 | -831 | -1373 | -1645 |
| 5 | mResp | ML | 1 | Intercept + Visit + TP (VC) | None | 137 | -1020 | -1565 | -1839 |
| 6 | mResp | ML | 1 | Intercept + Visit + TP (VC) | Visit (ID*TP) (AR1) | 138 | -1018 | -1567 | -1843 |
| *7 | mResp | ML | 1 | Intercept + Visit + TP (VC) | Visit (ID*TP) (CS) | 138 | 1553 | 1004 | 728 |
| *8 | mResp | ML | 1 | Intercept + Visit + TP (VC) | Visit (ID*TP) (SP_POWER) (t) | 138 | -1014 | -1563 | -1839 |
| ^9 | mResp | ML | 1 | Intercept + Visit + TP (VC) | Visit (ID*TP) (ARMA11) | 139 | -1008 | -1561 | -1839 |
| 10 | mResp | ML | 1 | Intercept + Visit + TP (VC) | Visit (ID*TP) (DIAG) | 147 | -970 | -1555 | -1849 |
| 11 | mResp | ML | 1 | Intercept + Visit + TP (VC) | Visit (ID*TP) (ARH1) | 148 | -967 | -1556 | -1852 |
| ^12 | mResp | ML | 1 | Intercept + Visit + TP (VC) | Visit (ID*TP) (CSH) | 148 | -966 | -1555 | -1851 |
| 13 | mResp | REML | 1 | Intercept + Visit + TP (VC) | Visit (ID*TP) (AR1) | 138 | -939 | -957 | -967 |
| Final | mResp | REML | 1 + $Iz_{between}$ $Iz_{between}$*Group $Iz_{between}$*Sex $Iz_{within}$ $Iz_{within}$*Group $Iz_{within}$*Sex | Intercept + Visit + TP (VC) | Visit (ID*TP) (AR1) | 144 | -909 | -927 | -937 |

*Failed to converge; ^Hessian matrix not positive definite; TP = resting assessment time point (Pre, Stim, Post)

| Model | Data | Method | Fixed (FE) | Random (Cov) | Repeated (Cov) | Param | BIC | AIC | -2 LL |
|---|---|---|---|---|---|---|---|---|---|
| 1 | sBP | ML | Age Group Sex Visit TP Visit*TP Group*Sex Group*Visit Group*Sex*Visit Group*TP Group*Sex*TP Group*Sex*Visit*TP | None | None | 90 | 2339 | 2017 | 1837 |
| 2 | sBP | ML | 1 | Intercept (VC) | None | 91 | 2127 | 1801 | 1619 |
| 3 | sBP | ML | 1 | Intercept + Visit (VC) | None | 92 | 2092 | 1763 | 1579 |
| *^4 | sBP | ML | 1 | Intercept + TP (VC) | None | 92 | 2132 | 1803 | 1619 |
| *^5 | sBP | ML | 1 | Intercept + Visit + TP (VC) | None | 93 | 2097 | 1765 | 1579 |
| 6 | sBP | ML | 1 | Intercept + Visit (VC) | Visit (ID*TP) (AR1) | 93 | 2096 | 1764 | 1578 |
| 7 | sBP | ML | 1 | Intercept + Visit (VC) | Visit (ID*TP) (CS) | 93 | 2087 | 1755 | 1569 |
| *8 | sBP | ML | 1 | Intercept + Visit (VC) | Visit (ID*TP) (SP_POWER) (t) | 93 | 2134 | 1802 | 1616 |
| *9 | sBP | ML | 1 | Intercept + Visit (VC) | Visit (ID*TP) (ARMA11) | 94 | 2102 | 1766 | 1578 |
| 10 | sBP | ML | 1 | Intercept + Visit (VC) | Visit (ID*TP) (DIAG) | 102 | 2129 | 1764 | 1560 |
| 11 | sBP | ML | 1 | Intercept + Visit (VC) | Visit (ID*TP) (ARH1) | 103 | 2133 | 1765 | 1559 |
| 12 | sBP | ML | 1 | Intercept + Visit (VC) | Visit (ID*TP) (CSH) | 103 | 2122 | 1754 | 1548 |
| 13 | sBP | REML | 1 | Intercept + Visit (VC) | Visit (ID*TP) (CS) | 93 | 1234 | 1221 | 1213 |
| Final | sBP | REML | 1 + $Iz_{between}$ $Iz_{between}$*Group $Iz_{between}$*Sex $Iz_{within}$ $Iz_{within}$*Group $Iz_{within}$*Sex | Intercept + Visit (VC) | Visit (ID*TP) (CS) | 99 | 1202 | 1190 | 1182 |

*Failed to converge; ^Hessian matrix not positive definite; TP = resting assessment time point (Pre, Stim, Post)

| Model | Data | Method | Fixed (FE) | Random (Cov) | Repeated (Cov) | Param | BIC | AIC | -2 LL |
|---|---|---|---|---|---|---|---|---|---|
| 1 | dBP | ML | Age Group Sex Visit TP Visit*TP Group*Sex Group*Visit Group*Sex*Visit Group*TP Group*Sex*TP Group*Sex*Visit*TP | None | None | 90 | 2195 | 1873 | 1693 |
| 2 | dBP | ML | 1 | Intercept (VC) | None | 91 | 2048 | 1723 | 1541 |
| 3 | dBP | ML | 1 | Intercept + Visit (VC) | None | 92 | 2008 | 1679 | 1495 |
| *^4 | dBP | ML | 1 | Intercept + TP (VC) | None | 92 | 2054 | 1725 | 1541 |
| 5 | dBP | ML | 1 | Intercept + Visit + TP (VC) | None | 93 | 2009 | 1676 | 1490 |
| 6 | dBP | ML | 1 | Intercept + Visit (VC) | Visit (ID*TP) (AR1) | 93 | 2013 | 1681 | 1495 |
| 7 | dBP | ML | 1 | Intercept + Visit (VC) | Visit (ID*TP) (CS) | 93 | 2009 | 1676 | 1490 |
| 8 | dBP | ML | 1 | Intercept + Visit (VC) | Visit (ID*TP) (SP_POWER) (t) | 93 | 2013 | 1681 | 1495 |
| *9 | dBP | ML | 1 | Intercept + Visit (VC) | Visit (ID*TP) (ARMA11) | 94 | 2014 | 1678 | 1490 |
| 10 | dBP | ML | 1 | Intercept + Visit (VC) | Visit (ID*TP) (DIAG) | 102 | 2036 | 1671 | 1467 |
| 11 | dBP | ML | 1 | Intercept + Visit (VC) | Visit (ID*TP) (ARH1) | 103 | 2041 | 1673 | 1467 |
| 12 | dBP | ML | 1 | Intercept + Visit (VC) | Visit (ID*TP) (CSH) | 103 | 2037 | 1669 | 1463 |
| 13 | dBP | REML | 1 | Intercept + Visit (VC) | Visit (ID*TP) (CSH) | 103 | 1215 | 1171 | 1143 |
| Final | dBP | REML | 1 + $Iz_{between}$ $Iz_{between}$*Group $Iz_{between}$*Sex $Iz_{within}$ $Iz_{within}$*Group $Iz_{within}$*Sex | Intercept + Visit (VC) | Visit (ID*TP) (CSH) | 109 | 1187 | 1143 | 1115 |

*Failed to converge; ^Hessian matrix not positive definite; TP = resting assessment time point (Pre, Stim, Post)

### E. Resting Phasic Physiological measures

| Model | Data | Method | Fixed (FE) | Random (Cov) | Repeated (Cov) | Param | BIC | AIC | -2 LL |
|---|---|---|---|---|---|---|---|---|---|
| 1 | SEPR | ML | Age Group Sex Visit Group*Sex Group*Visit Group*Sex*Visit | None | None | 46 | 344 | 211 | 119 |
| 2 | SEPR | ML | 1 | Intercept (VC) | None | 47 | 340 | 204 | 110 |
| ^3 | SEPR | ML | 1 | Intercept + Visit (VC) | None | 48 | 345 | 206 | 110 |
| 4 | SEPR | ML | 1 | Intercept (VC) | Visit (ID) (AR1) | 48 | 345 | 206 | 110 |
| *5 | SEPR | ML | 1 | Intercept (VC) | Visit (ID) (CS) | 48 | 361 | 223 | 127 |
| *6 | SEPR | ML | 1 | Intercept (VC) | Visit (ID) (SP_POWER) (t) | 48 | 368 | 230 | 134 |
| 7 | SEPR | ML | 1 | Intercept (VC) | Visit (ID) (ARMA11) | 49 | 348 | 207 | 109 |
| 8 | SEPR | ML | 1 | Intercept (VC) | Visit (ID) (DIAG) | 57 | 375 | 211 | 97 |
| 9 | SEPR | ML | 1 | Intercept (VC) | Visit (ID) (ARH1) | 58 | 380 | 213 | 97 |
| *10 | SEPR | ML | 1 | Intercept (VC) | Visit (ID) (CSH) | 58 | 388 | 221 | 105 |
| 11 | SEPR | REML | 1 | Intercept (VC) | None | 47 | 173 | 168 | 164 |
| *Final | SEPR | REML | 1 + $Iz_{between}$ $Iz_{between}$*Group $Iz_{between}$*Sex $Iz_{within}$ $Iz_{within}$*Group $Iz_{within}$*Sex | Intercept (VC) | None | 53 | 174 | 170 | 166 |

*Failed to converge; ^Hessian matrix not positive definite

| Model | Data | Method | Fixed (FE) | Random (Cov) | Repeated (Cov) | Param | BIC | AIC | -2 LL |
|---|---|---|---|---|---|---|---|---|---|
| 1 | SCR | ML | Age Group Sex Visit Group*Sex Group*Visit Group*Sex*Visit | None | None | 46 | 934 | 801 | 709 |
| 2 | SCR | ML | 1 | Intercept (VC) | None | 47 | 908 | 772 | 678 |
| ^3 | SCR | ML | 1 | Intercept + Visit (VC) | None | 48 | 912 | 774 | 678 |
| 4 | SCR | ML | 1 | Intercept (VC) | Visit (ID) (AR1) | 48 | 911 | 772 | 676 |
| *5 | SCR | ML | 1 | Intercept (VC) | Visit (ID) (CS) | 48 | 912 | 774 | 678 |
| *6 | SCR | ML | 1 | Intercept (VC) | Visit (ID) (SP_POWER) (t) | 48 | 919 | 780 | 684 |
| 7 | SCR | ML | 1 | Intercept (VC) | Visit (ID) (ARMA11) | 49 | 915 | 774 | 676 |
| 8 | SCR | ML | 1 | Intercept (VC) | Visit (ID) (DIAG) | 57 | 916 | 751 | 637 |
| 9 | SCR | ML | 1 | Intercept (VC) | Visit (ID) (ARH1) | 58 | 920 | 753 | 637 |
| *10 | SCR | ML | 1 | Intercept (VC) | Visit (ID) (CSH) | 58 | 920 | 753 | 637 |
| 11 | SCR | REML | 1 | Intercept (VC) | Visit (ID) (DIAG) | 57 | 564 | 534 | 510 |
| Final | SCR | REML | 1 + $Iz_{between}$ $Iz_{between}$*Group $Iz_{between}$*Sex $Iz_{within}$ $Iz_{within}$*Group $Iz_{within}$*Sex | Intercept (VC) | Visit (ID) (DIAG) | 63 | 543 | 514 | 490 |

*Failed to converge; ^Hessian matrix not positive definite

| Model | Data | Method | Fixed (FE) | Random (Cov) | Repeated (Cov) | Param | BIC | AIC | -2 LL |
|---|---|---|---|---|---|---|---|---|---|
| 1 | SEmRR | ML | Age Group Sex Visit Group*Sex Group*Visit Group*Sex*Visit | None | None | 46 | 1537 | 1404 | 1312 |
| 2 | SEmRR | ML | 1 | Intercept (VC) | None | 47 | 1523 | 1387 | 1293 |
| ^3 | SEmRR | ML | 1 | Intercept + Visit (VC) | None | 48 | 1528 | 1389 | 1293 |
| 4 | SEmRR | ML | 1 | Intercept (VC) | Visit (ID) (AR1) | 48 | 1527 | 1388 | 1292 |
| ^5 | SEmRR | ML | 1 | Intercept (VC) | Visit (ID) (CS) | 48 | 1528 | 1389 | 1293 |
| 6 | SEmRR | ML | 1 | Intercept (VC) | Visit (ID) (SP_POWER) (t) | 48 | 1526 | 1388 | 1292 |
| 7 | SEmRR | ML | 1 | Intercept (VC) | Visit (ID) (ARMA11) | 49 | 1533 | 1391 | 1293 |
| 8 | SEmRR | ML | 1 | Intercept (VC) | Visit (ID) (DIAG) | 57 | 1544 | 1380 | 1266 |
| 9 | SEmRR | ML | 1 | Intercept (VC) | Visit (ID) (ARH1) | 58 | 1549 | 1382 | 1266 |
| *10 | SEmRR | ML | 1 | Intercept (VC) | Visit (ID) (CSH) | 58 | 1360 | 1193 | 1077 |
| 11 | SEmRR | REML | 1 | Intercept (VC) | Visit (ID) (DIAG) | 57 | 978 | 949 | 925 |
| Final | SEmRR | REML | 1 + $Iz_{between}$ $Iz_{between}$*Group $Iz_{between}$*Sex $Iz_{within}$ $Iz_{within}$*Group $Iz_{within}$*Sex | Intercept (VC) | Visit (ID) (DIAG) | 63 | 933 | 904 | 880 |

*Failed to converge; ^Hessian matrix not positive definite

**Supplementary Table 4. Stimulation Intensities**

| Visit | CON | | | | | | VNS | | | | | |
|---|---|---|---|---|---|---|---|---|---|---|---|---|
| | Men | | | Women | | | Men | | | Women | | |
| 1 | 5.0 | ± | 0.9 | 3.8 | ± | 1.1 | 4.3 | ± | 1.6 | 2.6 | ± | 1.2 |
| 2 | 5.4 | ± | 1.0 | 4.4 | ± | 1.4 | 3.9 | ± | 0.9 | 3.3 | ± | 2.0 |
| 3 | 6.4 | ± | 1.4 | 4.7 | ± | 1.6 | 3.5 | ± | 1.1 | 2.9 | ± | 2.1 |
| 4 | 6.0 | ± | 2.5 | 4.0 | ± | 0.3 | 4.0 | ± | 0.3 | 3.0 | ± | 1.2 |
| 5 | 6.3 | ± | 1.7 | 3.1 | ± | 2.0 | 5.6 | ± | 0.7 | 2.8 | ± | 1.6 |
| 6 | 7.0 | ± | 2.0 | 4.5 | ± | 1.3 | 3.5 | ± | 1.5 | 4.3 | ± | 0.4 |
| 7 | 6.7 | ± | 2.5 | 3.9 | ± | 0.9 | 3.4 | ± | 1.8 | 3.0 | ± | 1.6 |
| 8 | 5.6 | ± | 1.2 | 4.5 | ± | 1.4 | 3.9 | ± | 2.3 | 2.4 | ± | 0.9 |
| 9 | 5.1 | ± | 1.8 | 4.9 | ± | 1.9 | 5.0 | ± | 3.3 | 2.9 | ± | 1.4 |
| 10 | 4.6 | ± | 1.3 | 4.3 | ± | 0.5 | 3.8 | ± | 2.3 | 4.1 | ± | 0.2 |
| F | 4.6 | ± | 1.9 | 3.9 | ± | 0.8 | 4.0 | ± | 1.0 | 4.8 | ± | 1.4 |
| **Group (*p*)** | **0.0473*** | | | | | | | | | | | |
| **Sex (*p*)** | **0.0598*** | | | | | | | | | | | |
| **Visit (*p*)** | **0.8220** | | | | | | | | | | | |
| **Group x Sex (*p*)** | **0.5133** | | | | | | | | | | | |
| **Group x Visit (*p*)** | **0.4149** | | | | | | | | | | | |
| **Sex x Visit (*p*)** | **0.3449** | | | | | | | | | | | |
| **Group x Sex X Visit (*p*)** | **0.5125** | | | | | | | | | | | |

Values are means ± SD (mA). Differences ($p \leq 0.10$) are indicated by asterisks after three-way ANOVA.
