## Supplemental Results for "Cognitive and autonomic physiological responses to daily transcutaneous auricular vagus nerve stimulation in healthy adults"

**S Results**

**Flanker.** The association between intensity and RT differed by sex (I$z_{between}$ x sex: $F_{1,4}$ = 45.17, $p$ = 0.0026). Women were faster than men at lower intensity (I$z_{between}$ = -1: MD = -126.3 ms [-198.1, -54.6], $t$(3.99) = -4.89, $p$ = 0.0081, $R_p^2$ = 0.857), similar at mean intensity (0: MD = 0.8 ms [-32.6, 34.2], $t$(3.99) = 0.07, $p$ = 0.9499, $R_p^2$ = 0.001), and slower at higher intensity (+1: MD = 127.9 ms [77.0, 178.9], $t$(3.99) = 6.98, $p$ = 0.0022, $R_p^2$ = 0.924). The association between intensity and rejected trials differed by group (I$z_{between}$ x group: $F_{1,21}$ = 6.77, $p$ = 0.0165). Individuals in VNS had more rejected trials than CON at lower intensity (I$z_{between}$ = -1: MD = 2.9 [0.6, 5.1], $t$(21.48) = 2.65, $p$ = 0.0148, $R_p^2$ = 0.246), did not differ at mean intensity (0: MD = -0.3 [-1.3, 0.6], $t$(21.73) = -0.69, $p$ = 0.4973, $R_p^2$ = 0.021), and had fewer rejected trials at higher intensity (+1: MD = -3.5 [-6.6, -0.4], $t$(21.43) = -2.34, $p$ = 0.0292, $R_p^2$ = 0.227). After adjusting for stimulation intensity, sex-specific differences between groups became more pronounced.

**WCST.** The association between intensity and RT differed by sex (I$z_{between}$ x sex: $F_{1,4}$ = 8.54, $p$ = 0.0447). Women were faster than men at lower intensity (I$z_{between}$ = -1: MD = -329.7 ms [-759.7, -100.4], $t$(3.89) = -2.15, $p_{adj}$ = 0.0997, $R_p^2$ = 0.543), similar at mean intensity (0: MD = -1.9 ms [-201.5, 197.6], $t$(3.92) = -0.03, $p_{adj}$ = 0.9793, $R_p^2$ = 0.000), and slower at higher intensity (+1: MD = 325.7 ms [20.5, 630.9], $t$(3.89) = 2.99, $p_{adj}$ = 0.0416, $R_p^2$ = 0.697). Adjustment for stimulation intensity eliminated differences in RT between men and women in VNS (MD = -182.5 ms [-512.0, 147.0], $t$(3.91) = -1.55, $p_{adj}$ = 0.1972, $R_p^2$ = 0.381; no adjustment: MD = -223.4 ms [-421.3, -25.4], $t$(7.09) = -2.66, $p_{adj}$ = 0.0320, $R_p^2$ = 0.499). Adjustment for intensity did not alter group/sex/visit effects on trial rejections.

**SWMT.** Adding stimulation intensity did not alter group/sex/visit effects for RT. The association between session-to-session variation in intensity and rejected trials differed by sex (I$z_{within}$ x sex: $F_{1,106}$ = 3.70, $p$ = 0.0569). Women had fewer rejected trials than men at lower intensity (I$z_{within}$ = -1: MD = -0.8 [-1.6, 0.0], $t$(26.96) = 1.95, $p_{adj}$ = 0.0623, $R_p^2$ = 0.123), and did not differ at mean (0: MD = -0.5 [-1.3, 0.3], $t$(20.49) = 1.40, $p_{adj}$ = 0.1786, $R_p^2$ = 0.087) or higher intensity (+1: MD = -0.3 [-1.1, 0.5], $t$(25.36) = 0.67, $p_{adj}$ = 0.5101, $R_p^2$ = 0.017). After adjusting for stimulation intensity, sex-specific between-group differences in trial rejections became more pronounced. Adjustment for intensity did not alter group/sex/visit effects on SWMT pupil size.

**Resting tonic autonomic physiological activity**. Tonic pupil size was unaffected by the inclusion of covariates for intensity. SCL was sensitive to differences in stimulation intensity between groups (I$z_{between}$ x group: $F_{1,4}$ = 7.17, $p$ = 0.0553), and sexes (I$z_{between}$ x sex: $F_{1,4}$ = 5.84, $p$ = 0.0731). Compared with CON, VNS had similar SCL at lower intensity (I$z_{between}$ = -1: MD =

-10.45 μS [-25.22, 4.33], $t(4) = -1.96$, $p_{adj} = 0.1211$, $R^2_p = 0.491$), and greater SCL at mean (0: MD = 5.68 μS [-0.55, 11.91], $t(4) = 2.53$, $p_{adj} = 0.0645$, $R^2_p = 0.616$) and higher intensity (+1: MD = 21.81 μS [1.35, 42.26], $t(4) = 2.96$, $p_{adj} = 0.0416$, $R^2_p = 0.687$). Women had higher SCL than men at lower intensity ($Iz_{between}$ = -1: MD = 13.21 μS [-3.06, 29.48], $t(4) = 2.25$, $p_{adj} = 0.0872$, $R^2_p = 0.559$), and similar SCL at mean intensity (0: MD = 2.85 μS [-4.72, 10.41], $t(4) = 1.05$, $p_{adj} = 0.3548$, $R^2_p = 0.215$) and higher intensity (+1: MD = -7.51 μS [-19.06, 4.03], $t(4) = -1.81$, $p_{adj} = 0.1451$, $R^2_p = 0.449$).

The relationship between intensity and mRR differed by sex ($Iz_{between}$ x sex: $F_{1, 4} = 4.92$, $p = 0.0908$). Women had shorter mRR than men at lower intensity ($Iz_{between}$ = MD = -1: -480.9 ms [-1051.8, 89.8], $t(4) = -2.34$, $p_{adj} = 0.0794$, $R^2_p = 0.578$), and equivalent mRR at mean intensity (0: MD = -147.3 ms [-412.6, 118.1], $t(4) = -1.54$, $p_{adj} = 0.1982$, $R^2_p = 0.372$) and higher intensity (+1: MD = 186.4 ms [-218.6, 591.5], $t(4) = 1.28$, $p_{adj} = 0.2730$, $R^2_p = 0.290$).

**Resting phasic autonomic physiological activity**. SEPR was directly related to stimulation intensity ($Iz_{between}$: $F_{1, 81} = 5.58$, $p = 0.0205$; -1: 0.65 mm [0.47, 0.83]; 0: 0.92 mm [0.78, 1.1]; +1: 1.19 mm [0.86, 1.52]). Treatment- and sex-specific differences in SEPR were eliminated when intensity was not included in the analysis (group: $F_{1, 7} = 3.19$, $p = 0.1175$; sex: $F_{1, 7} = 0.00$, $p = 0.9850$). SCR varied as a function of stimulation intensity and group ($Iz_{between}$ x group: $F_{1, 4} = 4.55$, $p = 0.1037$) as well as sex ($Iz_{between}$ x sex: $F_{1, 4} = 6.46$, $p = 0.0675$). Compared with CON, individuals in VNS had similar SCR at lower intensity ($Iz_{between}$ = -1: MD = -7.13 μS [-20.84, 6.59], $t(3.85) = -1.46$, $p = 0.2195$, $R^2_p = 0.358$), and greater SCR at mean ($Iz_{between}$ = 0) and higher intensity (+1: MD = 16.26 μS [-2.73, 35.25], $t(3.85) = 2.42$, $p_{adj} = 0.0756$, $R^2_p = 0.605$). In addition, women had higher SCR than men at lower intensity ($Iz_{between}$ = -1: MD = 13.40 μS [-1.69, 28.49], $t(3.84) = 2.51$, $p_{adj} = 0.0689$, $R^2_p = 0.621$), and similar SCR at mean intensity (0: MD = 3.49 μS [-3.46, 10.43], $t(4.05) = 1.39$, $p_{adj} = 0.2370$, $R^2_p = 0.322$) and higher intensity (+1: MD = -6.43 μS [-17.12, 4.27], $t(3.89) = -1.69$, $p_{adj} = 0.1687$, $R^2_p = 0.423$).
